# Multi-ancestry GWAS of major depression aids locus discovery, fine-mapping, gene prioritisation, and causal inference

**DOI:** 10.1101/2022.07.20.500802

**Authors:** Xiangrui Meng, Georgina Navoly, Olga Giannakopoulou, Daniel Levey, Dora Koller, Gita Pathak, Nastassja Koen, Kuang Lin, Miguel E. Rentería, Yanzhe Feng, J. Michael Gaziano, Dan J. Stein, Heather J. Zar, Megan L. Campbell, David A. van Heel, Bhavi Trivedi, Sarah Finer, Andrew McQuillin, Nick Bass, V. Kartik Chundru, Hilary Martin, Qin Qin Huang, Maria Valkovskaya, Po-Hsiu Kuo, Hsi-Chung Chen, Shih-Jen Tsai, Yu-Li Liu, Kenneth S. Kendler, Roseann E. Peterson, Na Cai, Yu Fang, Srijan Sen, Laura Scott, Margit Burmeister, Ruth Loos, Michael Preuss, Ky’Era V. Actkins, Lea K. Davis, Monica Uddin, Agaz Wani, Derek Wildman, Robert J. Ursano, Ronald C. Kessler, Masahiro Kanai, Yukinori Okada, Saori Sakaue, Jill Rabinowitz, Brion Maher, George Uhl, William Eaton, Carlos S. Cruz-Fuentes, Gabriela A. Martinez-Levy, Adrian I. Campos, Iona Y. Millwood, Zhengming Chen, Liming Li, Sylvia Wassertheil-Smoller, Yunxuan Jiang, Chao Tian, Nicholas G. Martin, Brittany L. Mitchell, Enda M. Byrne, Naomi R. Wray, Swapnil Awasthi, Jonathan R. I. Coleman, Stephan Ripke, PGC MDD Working Group, China Kadoorie Biobank Collaborative Group, the 23andMe Research Team, Genes & Health Research Team, Tamar Sofer, Robin G. Walters, Renato Polimanti, Erin C. Dunn, Murray B. Stein, Joel Gelernter, Cathryn Lewis, Karoline Kuchenbaecker

## Abstract

Most genome-wide association studies (GWAS) of major depression (MD) have been conducted in samples of European ancestry. Here we report a multi-ancestry GWAS of MD, adding data from 21 studies with 88,316 MD cases and 902,757 controls to previously reported data from individuals of European ancestry. This includes samples of African (36% of effective sample size), East Asian (26%) and South Asian (6%) ancestry and Hispanic/Latinx participants (32%). The multi-ancestry GWAS identified 190 significantly associated loci, 53 of them novel. For previously reported loci from GWAS in European ancestry the power-adjusted transferability ratio was 0.6 in the Hispanic/Latinx group and 0.3 in each of the other groups. Fine-mapping benefited from additional sample diversity: the number of credible sets with ≤5 variants increased from 3 to 12. A transcriptome-wide association study identified 354 significantly associated genes, 205 of them novel. Mendelian Randomisation showed a bidirectional relationship with BMI exclusively in samples of European ancestry. This first multi-ancestry GWAS of MD demonstrates the importance of large diverse samples for the identification of target genes and putative mechanisms.

## Background

Major depression (MD) is one of the most pressing global health challenges^1^. It affects an estimated 300 million people, 4.4% of the world’s population^2^. Depression has been linked to a wide range of adverse health outcomes, including cardiovascular disease, asthma, cancer, poor maternal and child health, substance abuse and suicide, and it also affects family relationships and work performance^3–5^.

Molecular genetic research has shown promise of uncovering biological mechanisms underlying the development of MD^6,7^. However, previous genome-wide association studies (GWAS) revealed that the genetic architecture of MD is highly polygenic, characterised by variants that individually confer small risk increases^8^. This creates challenges for identification of variants that reproducibly provide genome-wide levels of association. The heterogeneity of MD symptoms may be mirrored by aetiological heterogeneity and that has been highlighted as a likely reason for the small genetic effects^9^. Therefore, previous genetic research explored the impact of different outcome definitions^6,10^, sex^11–13^ and trauma exposure^14–16^ on heterogeneity. However, the role of ancestry and ethnicity in the genetics of MD has not yet been systematically evaluated.

Previous GWAS of MD were mostly conducted in individuals of European ancestry^6,17–19^. The largest MD GWAS to date, combined data from several studies, including the Million Veteran Program (MVP), the Psychiatric Genomics Consortium Major Depressive Disorder group (PGC-MDD), the UK Biobank, FinnGen and 23andMe, resulting in a total of 1,154,267 participants of European ancestry (340,591 cases) and identifying 223 independent significant single nucleotide variants (SNVs) at 178 genomic loci linked to MD^19^. The study also included data from 59,600 African Americans (25,843 cases) from the MVP cohort. In their bi-ancestral meta-analysis, the number of significant SNVs increased to 233. Other MD GWASs were conducted in African American and Hispanic/Latinx participants with limited sample sizes and did not find variants with statistically significant associations with MD^20,21^.

With 10,640 female Chinese participants, the CONVERGE study is the largest MD GWAS conducted outside “Western” countries to date^7^. The study identified two genome-wide significant associations linked to mitochondrial biology and reported a genetic correlation of 0.33 with MD in European ancestry samples^22^. In line with this, our recent work demonstrated that some of the previously identified loci from GWAS conducted in samples of European ancestry are not transferable to samples of East Asian ancestry^23^. The analyses also revealed differences in genetic associations by geographic region (East Asian versus European/American countries) for ancestry-matched participants. These findings suggest that there can be heterogeneity in genetic risk factors for MD by ancestry as well as cultural context.

This heterogeneity could impact on findings of genetic studies when evaluating causal effects of risk factors for MD. In particular, previous studies in samples of European ancestry reported genetic correlations and causal relationships between MD and cardiometabolic outcomes^6,17,19,24^. Notably, our previous study indicated a contradicting direction for associations between MD and BMI in East Asian individuals (negative genetic correlation) and European ancestry individuals (positive genetic correlation)^23^. Thus, investigating causal relationships in diverse ancestry groups and in different disease subtypes is important to ensure generalisability and to distinguish between biological and societal mechanisms underlying the relationship between a risk factor and the disease.

Increasing diversity in genetic research is also important to ensure equitable health benefits^25^. In the United States of America, differences in presentation of MD across ethnic groups can impact on the likelihood of diagnosis^26^. Genetics optimised for European ancestry participants would primarily benefit that group of patients and could therefore further widen the disparities in diagnosis and treatment between groups.

We used data from samples with diverse ancestry and carried out genome-wide association meta-analyses, followed by fine-mapping and prioritisation of target genes. We assessed transferability of genetic loci across ancestry groups. Finally, we explored bi-directional causal links between MD and cardiometabolic traits.

## Results

### GWAS in African, East Asian, South Asian, and Hispanic/Latinx samples

We first conducted GWAS meta-analyses stratified by ancestry/ethnic group. Individuals were assigned to ancestry groups (African, South Asian, East Asian, European) using principal components analyses based on genetic relatedness matrices. We acknowledge the arbitrary nature of these groupings and the imprecision of this approach. However, creating such groups enabled us to look for associations that are specific to certain groups and to assess the transferability of previously identified loci. Assignment to the Hispanic/Latinx group was based on self-report or on recruitment in a Latin American country. The studies included in the meta-analyses used a range of measures to define MD: structured clinical interviews, medical healthcare records, symptoms questionnaires, and self-completed surveys.

The analyses included 36,818 MD cases and 161,679 controls of African ancestry (λ_1000_=1.001, Linkage disequilibrium score regression (LDSC) intercept = 1.034), 21,980 cases and 360,956 controls of East Asian ancestry (λ_1000_=1.002, LDSC intercept = 1.026), 4,505 cases and 27,176 controls of South Asian ancestry (λ_1000_=1.002, LDSC intercept = 1.014) as well as 25,013 cases and 352,946 controls in the Hispanic/Latinx group (λ_1000_=1.002, LDSC intercept = 1.051) (**Supplementary Figures 2-5**). To account for the minor inflation found in the Hispanic/Latinx samples, test statistics for this analysis were corrected based on the LDSC intercept.

In the Hispanic/Latinx group, the G-allele of rs78349146 at 2q24.2 was associated with increased risk of MD (effect allele frequency (EAF) = 0.04, beta = 0.15, standard error (SE) = 0.03, *P* value = 9.32×10^−9^) (**Supplementary Figures 1, 3**). The variant is located downstream of the gene *LINC01806* (long intergenic non-protein coding RNA 1806). Testing multi-ancestry brain expression quantitative trait loci (eQTLs)^27^, we observed significant colocalisation for *DPP4, RBMS1*, and *TANK*. We tested ancestry specific eQTLs from blood, and observed *RBMS1* (H3:PP (Hispanic/Latinx) = 99.12%). For the protein quantitative trait loci (pQTLs) from blood, we either did not find the genes or were not enough to test for colocalisation (**Supplementary Table 1**).

No variants were associated at genome-wide significance in the GWAS in samples of African, East and South Asian ancestry (**Supplementary Figures 2**,**4**,**5**). One locus was suggestively associated in the African ancestry GWAS (**Supplementary Figures 1-2**). The lead variant, rs6902879 (effect allele: A, EAF = 0.16, beta = -0.08, SE = 0.01, *P* value=5.3×10^−8^) at 6q16 is located upstream of the melanin-concentrating hormone receptor 2 gene (*MCHR2* and associated with increased expression of *MCHR2* in cortex based on Genotype-Tissue Expression (GTEx v8) (*P* = 6.0×10^−6^). Testing the multi-ancestry brain eQTLs^27^, we observed significant colocalisation for *GRIK2* and *ASCC3* with significant ancestry differences for *ASCC3* (H3:PP (European ancestry) = 99.97%). *MCHR2* was not present in the RNA data.

Although the lead variants at 2q24.2 and at 6q16 did not display strong evidence of association in a large published GWAS in participants of European ancestry^18^ (*P*>0.01), in each case there was an uncorrelated variant within 500Kb of the lead variants associated at *P* <10^−6^.

### Transferability of MD associations across ancestry groups

Previous GWAS in samples of European ancestry have identified 196 loci associated with MD^17–19^. We assessed whether these genetic associations are shared across different ancestry groups. Individual loci may be underpowered to demonstrate an association, therefore, we followed an approach we recently developed^28^ and first estimated the number of loci we expect to see an association for when accounting for sample size (N), linkage disequilibrium (LD), and minor allele frequency (MAF). This estimate varied widely between ancestry groups, e.g. we expected to detect significant associations for 65% of MD loci in the GWAS with samples of African ancestry but only for 15% of MD loci in samples of South Asian ancestry (**Figure 1, Supplementary Table 2**). We report the power-adjusted transferability (PAT) ratio, i.e. the observed number divided by the expected number of loci. Transferability was low, with PAT ratios of 0.27 in African ancestry samples, and 0.29 in both East Asian and South Asian ancestry samples. In the Hispanic/Latinx group, the PAT ratio was 0.63, notably higher than in the other groups. We also assessed the transferability of 102 loci identified in the PGC-MDD GWAS^17^ in an independent study in samples of European ancestry, the Australian Genetics of Depression Study (AGDS)^18^. The PAT ratio was 1.48, considerably higher than the cross-ancestry PAT estimates.

**Figure 1.**
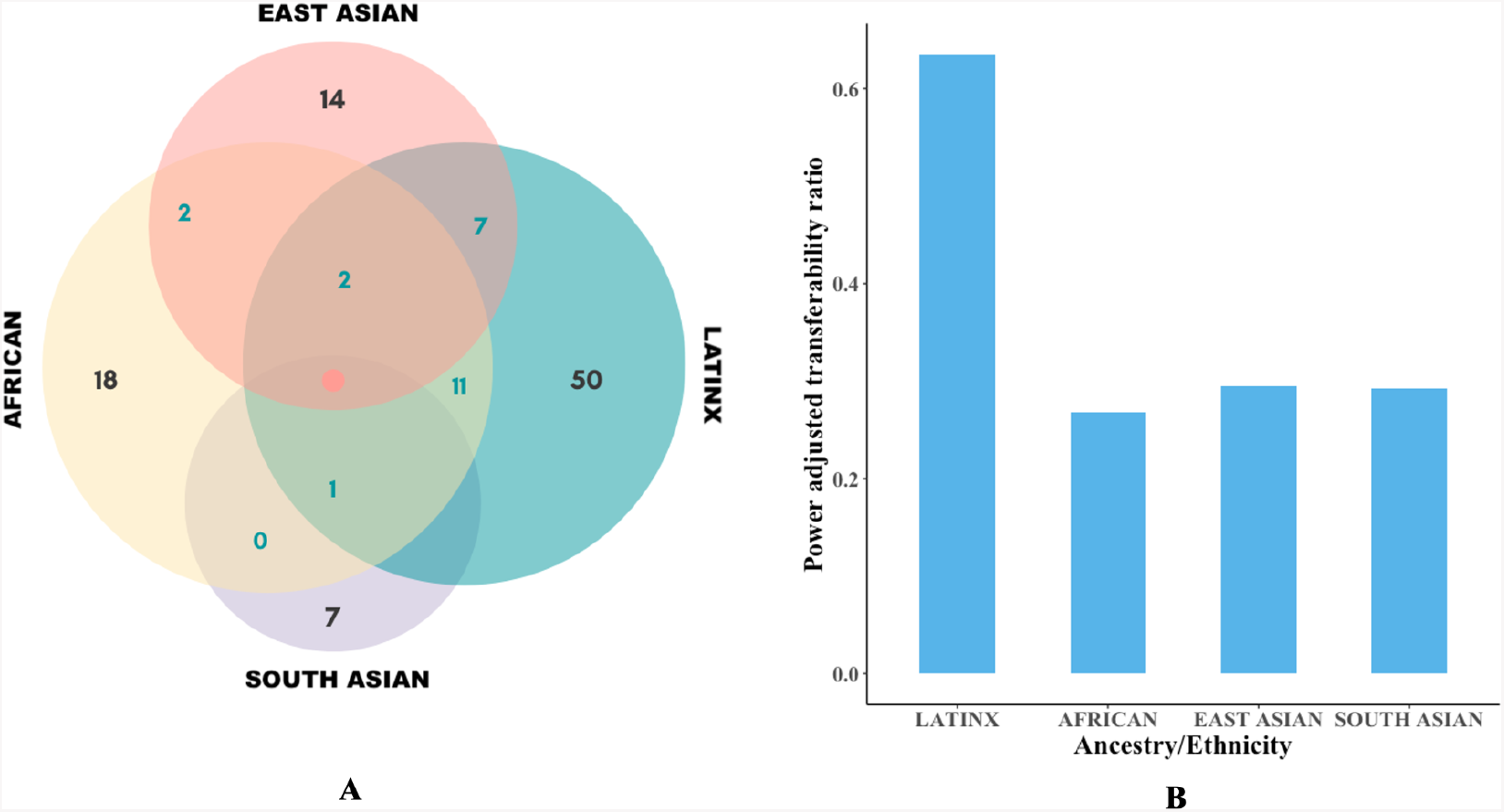
Transferability of previously reported loci from European ancestry discovery GWAS of major depression to other ancestry groups. Panel **(A)** shows the numbers of previously identified loci from European ancestry studies which were transferable to each of the other ancestry/ethnic groups: African, Hispanic/Latinx, South Asian and East Asian (in black) and their intersections (in cyan). Panel **(B)** shows Power Adjusted Transferability ratios, calculated from the observed number of transferable loci divided by the expected number of transferable loci, taking effect estimates from previous European ancestry studies, and allele frequency and sample size information from our African, Hispanic/Latinx, South Asian and East Asian ancestry cohorts.

### Multi-ancestry meta-analysis

We carried out a multi-ancestry meta-analysis using data from studies conducted in participants of African, East Asian and South Asian ancestry and Hispanic/Latinx samples (Supplementary material) and combined them with previously published data for 258,364 cases and 571,252 controls of European ancestry^17,18^, yielding a total sample size of 345,389 cases and 1,469,702 controls. These analyses provided results for 22,941,580 SNVs after quality control. There was no evidence of residual population stratification (λ_1000_=1.001, LDSC intercept = 1.019). We identified 190 independent genome-wide significant SNVs mapping to 169 loci that were separated from each other by at least 500kb (**Supplementary Table 3**). 53 of SNVs represent novel associations (r^2^ <0.1 and located > ±250kb from previously reported variants).

### Multi-ancestry fine-mapping

We used a multi-ancestry Bayesian fine-mapping method^29^ to derive 99% credible sets for 155 loci that were associated at genome-wide significance and did not show evidence of multiple independent signals. For comparison, we also implemented single ancestry fine-mapping of the same loci based on GWAS conducted in participants of European ancestry, including the PGC-MDD and the AGDS^17,18^.

The multi-ancestry fine-mapping increased fine-mapping resolution substantially as compared with fine-mapping solely based on the data from European ancestry participants. The median size of the 99% credible sets was reduced from 65.5 to 30 variants. Among the 145 loci for which we conducted fine-mapping on both sample sets, 113 (77.9%) loci had a smaller 99% credible set from the multi-ancestry fine-mapping, whilst 4 loci (0 from the European fine-mapping) were resolved to single putatively causal SNVs and 32 loci were resolved to no more than 10 putatively causal SNVs each (**Figure 2**).

**Figure 2.**
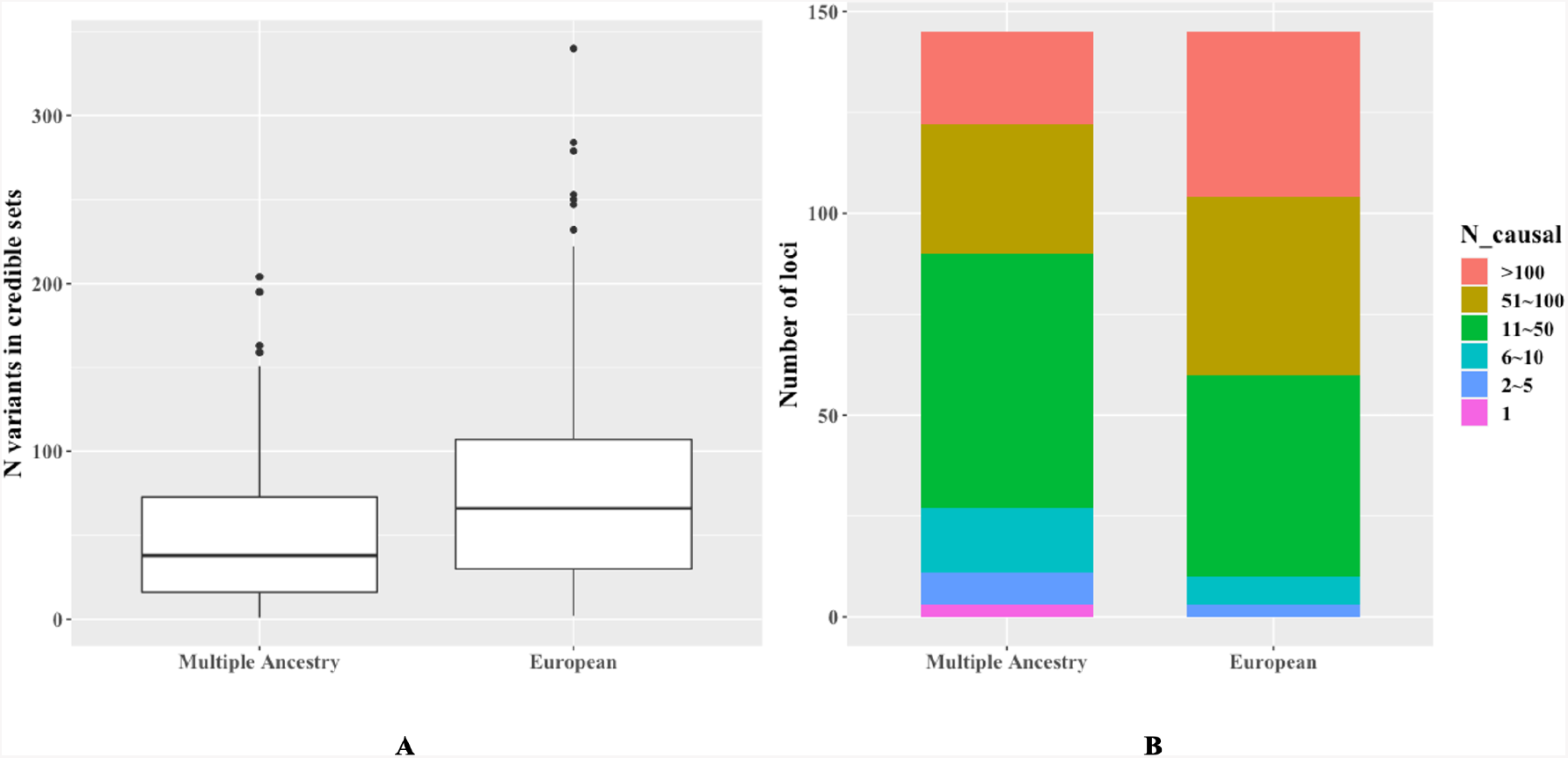
Resolution of the locus fine-mapping based on the multi-ancestry and the European ancestry GWAS, showing the size of the credible sets for 155 significant loci. A) box plot showing the median and interquartile range of credible set sizes. B) stacked bar charts showing the number of loci within size categories.

For several regions, we successfully fine-mapped the credible sets down to one variant (**Supplementary Table 4**). rs12699323 annotated as an intron variant and linked to the expression of gene Transmembrane Protein 106B *(TMEM106B)*. rs1806152 is a splice region variant, associated with the expression of the nearby gene PAX6 Upstream Antisense RNA *(PAUPAR)* on chromosome 11. Finally, rs9564313, annotated as an intron variant, has been linked to expression of *gene PCDH9* (Protocadherin-9); a gene that has also been highlighted as having an important function in our TWAS and MAGMA results^30,31^.

### Transcriptome-wide association study and gene prioritisation

To better understand the biological mechanisms of our GWAS findings, we performed several *in silico* analyses to functionally annotate and prioritise the most likely causal genes. We carried out a transcriptome-wide association study (TWAS) for expression in tissues relevant to MD^32^. We combined the TWAS results with Functional Mapping and Annotation (FUMA), conventional and HiC-MAGMA, to prioritise target genes. We also carried out ancestry-specific eQTL and pQTL colocalisation analyses **(Supplementary Table 5)**.

The TWAS identified 354 significant associations (*P* < 1.37×10^−6^) with MD, 205 of which had not been previously reported (**Figure 3, Supplementary Table 6**). The two most significant gene associations with MD were *RPL31P12* (GTEx Brain Cerebellum, Z = -10.68, *P* = 1.27×10^−26^) and *NEGR1* (GTEx Brain Caudate Basal Ganglia, Z = 10.677, *P* = 1.30×10^−26^). Both results are consistent with findings of a previous, large TWAS on MD in samples of European ancestry ^32^.

**Figure 3.**
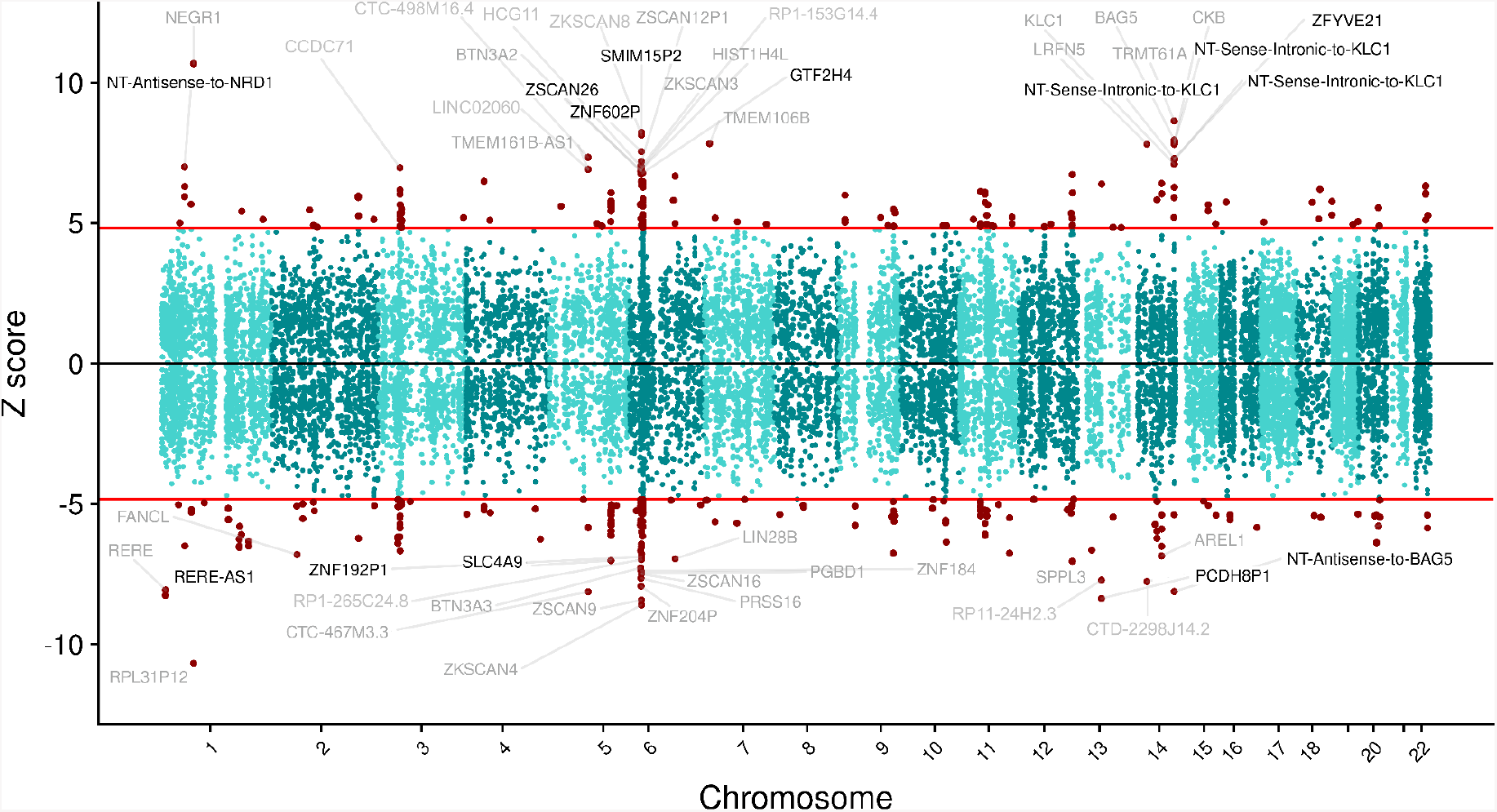
The relationship between gene expression and major depression. based on a multi-ancestry, multi-tissue transcriptome-wide association study (TWAS). Manhattan-style z-score plot of the TWAS, showing all tested genes as single dots across all autosomes. Significant gene associations are shown as red dots (354 significant genes, 205 of them novel) and the 50 most significant gene names are highlighted on both sides of the plot. Novel genes are shown in black, while genes previously associated with MD are shown in grey. The red lines indicate the transcriptome-wide significance threshold (*P*< 1.37×10-6). Genes on the top part of the graph have a positive direction of effect (direct effect), while the genes on the bottom part of the plot have a negative direction of effect (inverse effect). NT - Novel Transcript

Amongst the 205 novel gene associations with MD, genes *PCDH8P1* (GTEx Brain Anterior cingulate cortex BA24, Z = -8.3679, *P* = 5.86×10^−17^), *RERE-AS1* (GTEx Thyroid, Z = -8.26, *P* = 1.48×10^−16^) and *SMIM15P2* (GTEx Brain Cerebellar Hemisphere, Z = 8.2281, *P* = 1.90×10^−16^) were the most significant novel TWAS results, associated with MD. *NDUFAF3* was another novel gene association with MD (GTEx Brain Nucleus Accumbens Basal Ganglia, Z-score = -5.0785, *P* = 3.80×10^−7^, Best GWAS ID = rs7617480, Best GWAS *P* = 0.00001). These results are also confirmed by HiC-MAGMA. The protein *NDUFAF3* encodes is targeted by Metformin, the first-line drug for the treatment of type-2 diabetes.

43 genes displayed evidence of association across all the four gene prioritisation methods mentioned, and were classified as high confidence genes (**Table 1, Supplementary Tables 7-10**). These included genes repeatedly highlighted in previous studies due to their strong evidence of association and biological relevance: *NEGR1, DRD2, CELF4, LRFN5, TMEM161B, and TMEM106B*. 25 of the high confidence genes encode targets of existing drugs, such as Simvastatin (*RHOA*).

**Table 1.** Genes associated with major depression, including 12 genes significantly associated in the TWAS and Hi-C MAGMA, i.e., not in physical proximity to a GWAS hit, and 43 genes significantly associated across all four methods (TWAS, FUMA, MAGMA, Hi-C MAGMA).

| Gene<sup>a</sup> | Drug<sup>b</sup> | FUMA<sup>c</sup> | MAGMA<sup>d</sup> | Hi-C MAGMA<sup>d</sup> | TWAS P | Novel<sup>e</sup> | Credible set<sup>f</sup> |
| --- | --- | --- | --- | --- | --- | --- | --- |
| **Genes associated in TWAS and Hi-C MAGMA** |  |  |  |  |  |  |  |
| NDUFAF3 | Metformin, NADH | No | 1.00 | 0.004 | 3.80×10<sup>-7</sup> | Yes | 9 |
| PBRM1 | Alprazolam Durvalumab Everolimus | No | 0.10 | 0.017 | 3.20×10<sup>-7</sup> | Yes | - |
| TBCA | - | No | 0.16 | 0.042 | 1.29×10<sup>-6</sup> | Yes | - |
| BTN2A3P | - | No | 1.00 | 0.000 | 2.33×10<sup>-8</sup> | Yes | - |
| ZNF204P | - | No | 1.00 | 0.014 | 2.13×10<sup>-15</sup> | No | - |
| HLA-B | Thalidomide Ticlopidine Phenobarbital Carbamazepine Clozapine, Lamotrigine | No | 0.46 | 0.000 | 1.13×10<sup>-7</sup> | No | - |
| RABGAP1 | - | No | 1.00 | 0.001 | 1.91×10<sup>-8</sup> | Yes | - |
| GOLGA1 | - | No | 1.00 | 0.020 | 1.56×10<sup>-7</sup> | Yes | - |
| FRAT2 | - | No | 0.78 | 0.017 | 9.39×10<sup>-7</sup> | Yes | - |
| ENSG00000278376 | - | No | 0.06 | 0.004 | 6.55×10<sup>-7</sup> | - | 62 |
| TRHDE-AS1 | - | No | 0.14 | 0.048 | 6.92×10<sup>-7</sup> | Yes | - |
| INSYN1-AS1 | - | No | 0.25 | 0.014 | 5.53×10<sup>-8</sup> | Yes | 25 |
| **Genes associated across all four methods** |  |  |  |  |  |  |  |
| RERE | - | Yes | 0.000 | 0.000 | 7.35×10<sup>-16</sup> | No | 45 |
| NEGR1 | - | Yes | 0.000 | 0.000 | 1.30×10<sup>-26</sup> | No | - |
| ZNF638 | Cytidine | Yes | 0.003 | 0.004 | 5.64×10<sup>-7</sup> | No | 61 |
| RFTN2 | Lipopolysaccharide | Yes | 0.003 | 0.000 | 1.52×10<sup>-7</sup> | No | 204 |
| ZNF445 | - | Yes | 0.000 | 0.001 | 1.52×10<sup>-10</sup> | No | 138 |
| ZNF197 | - | Yes | 0.000 | 0.000 | 4.61×10<sup>-10</sup> | No | 138 |
| CCDC71 | - | Yes | 0.000 | 0.039 | 3.12×10<sup>-12</sup> | Yes | 9 |
| ENSG00000225399 | - | Yes | 0.010 | 0.003 | 1.07×10<sup>-8</sup> | - | 9 |
| RHOA | Simvastatin Pravastatin Atorvastatin Magnesium CCG-1423 | Yes | 0.031 | 0.019 | 1.45×10<sup>-7</sup> | No | 9 |
| CDH9 | Calcium | Yes | 0.003 | 0.002 | 2.17×10<sup>-8</sup> | No | 95 |
| TMEM161B | Crofelemer | Yes | 0.000 | 0.000 | 5.26×10<sup>-9</sup> | No | - |
| PFDN1 | - | Yes | 0.025 | 0.000 | 5.60×10<sup>-8</sup> | Yes | 67 |
| SLC4A9 | Sodium bicarbonate | Yes | 0.029 | 0.002 | 2.25×10<sup>-12</sup> | No | 67 |
| HARS1 | Adenosine phosphate, Pyrophosphate, Phosphate, Histidine | Yes | 0.024 | 0.017 | 5.29×10<sup>-8</sup> | Yes | 141 |
| HARS2 | Adenosine phosphate Pyrophosphate, Phosphate Histidine | Yes | 0.019 | 0.044 | 2.32×10<sup>-8</sup> | No | 141 |
| ZMAT2 | - | Yes | 0.014 | 0.005 | 1.11×10<sup>-9</sup> | No | 141 |
| PCDHA1 | Calcium | Yes | 0.015 | 0.005 | 1.15×10<sup>-8</sup> | No | 141 |
| PCDHA2 | Calcium | Yes | 0.031 | 0.010 | 1.55×10<sup>-8</sup> | No | 141 |
| PCDHA3 | Calcium | Yes | 0.043 | 0.004 | 1.06×10<sup>-8</sup> | No | 141 |
| TMEM106B | Crofelemer | Yes | 0.000 | 0.000 | 4.87×10<sup>-15</sup> | No | 1 |
| ZDHHC21 | Coenzyme A, Palmitoyl-CoA | Yes | 0.002 | 0.036 | 5.13×10<sup>-7</sup> | No | 42 |
| SORCS3 | - | Yes | 0.000 | 0.000 | 1.98×10<sup>-10</sup> | No | 16 |
| MYBPC3 | - | Yes | 0.004 | 0.012 | 9.23×10<sup>-10</sup> | No | 48 |
| SLC39A13 | Zinc chloride Zinc sulfate | Yes | 0.007 | 0.003 | 9.14×10<sup>-7</sup> | Yes | 48 |
| CTNND1 | - | Yes | 0.002 | 0.010 | 1.84×10<sup>-7</sup> | No | 60 |
| ANKK1 | Methadone, Naltrexone, Fostamatinib | Yes | 0.011 | 0.000 | 1.41×10<sup>-11</sup> | No | - |
| DRD2 | Cabergoline, Ropinirole, Sulpiride | Yes | 0.000 | 0.000 | 3.95×10<sup>-8</sup> | No | - |
| MLEC | - | Yes | 0.013 | 0.000 | 8.90×10<sup>-7</sup> | Yes | - |
| SPPL3 | - | Yes | 0.000 | 0.000 | 1.89×10<sup>-12</sup> | No | - |
| *LRFN5* | - | Yes | 0.000 | 0.000 | $5.79 \times 10^{-15}$ | No | 10 |
| *AREL1* | - | Yes | 0.007 | 0.000 | $7.24 \times 10^{-12}$ | No | 143 |
| *DLST* | Lipoic acid Succinyl-CoA, Coenzyme A, Dihydrolipoamide (S)--Succinyl-dihydrolipoamide | Yes | 0.000 | 0.001 | $1.51 \times 10^{-9}$ | No | 143 |
| *MARK3* | Fostamatinib, Alsterpaullone | Yes | 0.000 | 0.000 | $3.50 \times 10^{-9}$ | No | 11 |
| *KLC1* | Fluorouracil Irinotecan leucovorin | Yes | 0.000 | 0.000 | $1.26 \times 10^{-12}$ | No | 11 |
| *XRCC3* | Fluorouracil Irinotecan leucovorin | Yes | 0.004 | 0.000 | $3.49 \times 10^{-10}$ | No | 11 |
| *ZFYVE21* | Inositol 1,3-Bisphosphate | Yes | 0.000 | 0.000 | $2.83 \times 10^{-13}$ | Yes | 11 |
| *CELF4* | Iloperidone | Yes | 0.000 | 0.000 | $9.66 \times 10^{-9}$ | No | 8 |
| *RAB27B* | Guanosine-5'-Diphosphate | Yes | 0.000 | 0.042 | $5.61 \times 10^{-10}$ | No | - |
| *EMILIN3* | - | Yes | 0.039 | 0.001 | $6.66 \times 10^{-8}$ | No | 64 |
| *CHD6* | Phosphate, ATP, ADP | Yes | 0.001 | 0.001 | $1.76 \times 10^{-10}$ | No | 64 |
| *EP300* | Acetyl-CoA, TGF-beta, Garcinol cyclic amp, Curcumin, Mocetinostat | Yes | 0.000 | 0.000 | $1.71 \times 10^{-9}$ | No | 16 |
| *RANGAP1* | - | Yes | 0.000 | 0.003 | $6.73 \times 10^{-9}$ | No | 16 |
| *ZC3H7B* | - | Yes | 0.001 | 0.000 | $1.37 \times 10^{-9}$ | No | 16 |

<sup>a</sup>Ensembl IDs were shown for genes without symbol names. <sup>b</sup>Drugs targeting the prioritised genes or genes of the same family from GeneCards, DrugBank and ChEMBL. <sup>c</sup>Gene mapped by FUMA positional mapping or eQTL mapping. <sup>d</sup>Bonferroni adjusted *P* value for MAGMA or Hi-C MAGMA (*P* < 0.05 implies statistical significance). <sup>e</sup>Novel report as compared with previous MAGMA and TWAS on MD. <sup>f</sup>Number of variants in the 99% credible set for the mapped locus from multi ancestry fine mapping.

### Mendelian Randomisation

We assessed bi-directional causal relationships between MD and cardiometabolic traits using ancestry-specific two-sample Mendelian Randomisation analyses. Our results indicated a positive, bi-directional relationship between MD and BMI (MD->BMI: β = 0.092, 95% CI = 0.024-0.161, *P* = 8.12×10^−3^, BMI->MD: β = 0.138, 95% CI = 0.097-0.180, *P* = 6.88×10^−11^) (**Figure 4, Supplementary Table 11**). This bi-directional relationship was exclusively observed in samples of European ancestry (*P* > 0.1 in all other groups). MD was also causal for other indicators of unfavourable metabolic profiles in samples of European ancestry: triglycerides (TG, positive effect; β = 0.116, 95% CI = 0.070-0.162, *P* = 7.93×10^−7^), HDL (negative effect; β = -0.058, 95% CI = -0.111--0.006, *P* = 0.029), and LDL cholesterol (positive effect; β = 0.054, 95% CI = 0.012-0.096, *P* = 0.011). The effects remained significant after removing the variants contributing to the possible heterogeneity bias observed through the MR-PRESSO global test **(Supplementary Table 11B)**. In samples of East Asian ancestry, on the other hand, we found a negative causal association between TG and MD (β = -0.127, 95% CI = -0.223--0.032, *P* = 9.22×10^−3^). Moreover, MD showed a positive causal association with systolic blood pressure (SBP) (β = 0.034, 95% CI = 0.009-0.059, *P* = 7.66×10^−3^). In samples of African ancestry, SBP had a positive causal association with MD (β = 0.080, 95% CI = 0.026-0.133, *P* = 3.43×10^−3^).

**Figure 4.**
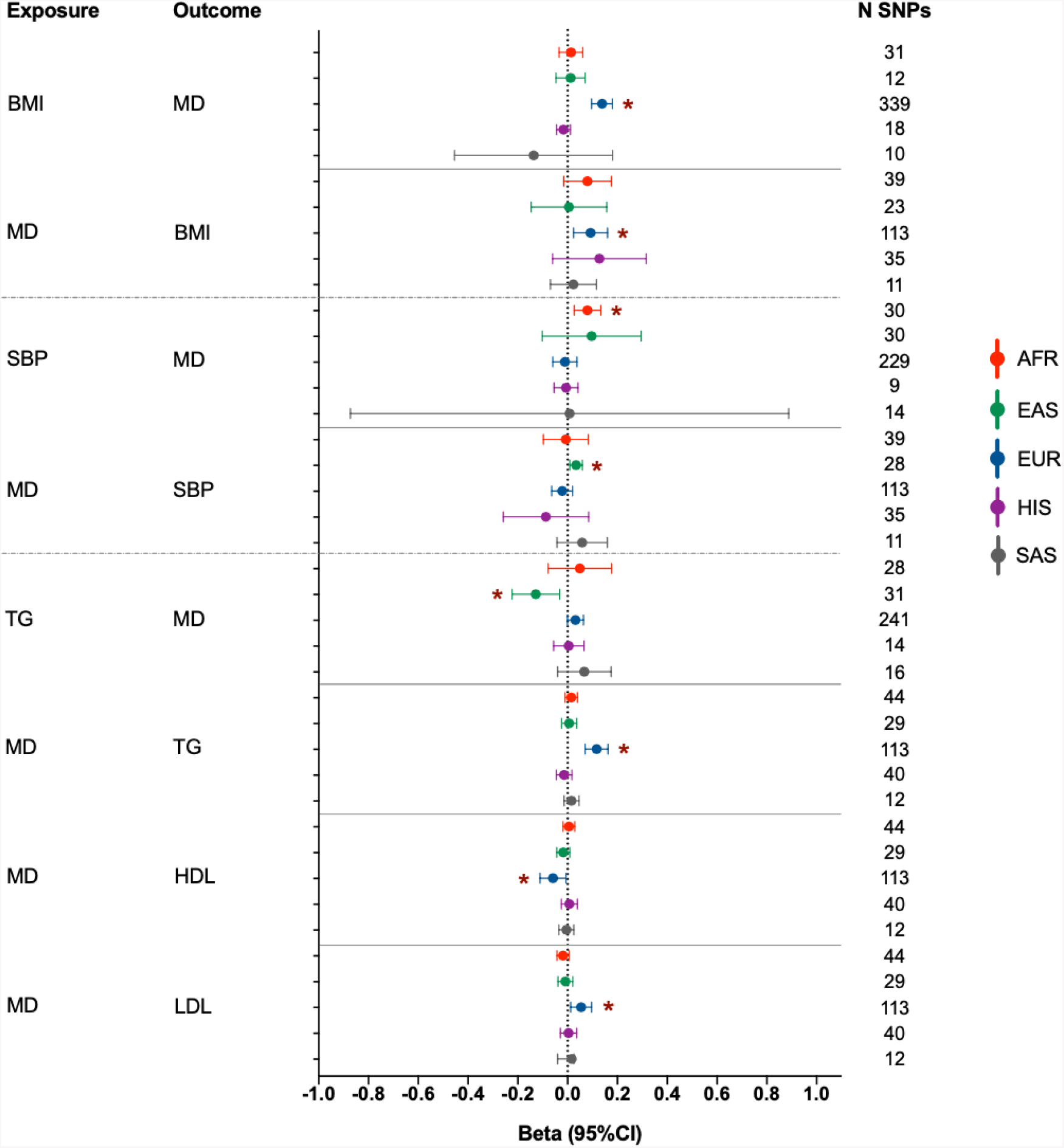
Bi-directional Mendelian Randomization analyses between major depression and cardiometabolic outcomes. The associations are shown with a beta and a 95% confidence interval. Significant associations are marked with a red asterisk. The number of SNPs refers to the number of variants used as instrumental variables in each analysis. Results not shown for diastolic blood pressure for which there were no significant associations. Abbreviations: AFR: African, EAS: East Asian, EUR: European, HIS: Hispanic/Latinx, SAS: South Asian, BMI: body mass index, MD: major depression, TG: triglycerides, SBP: systolic blood pressure, HDL: high-density lipoprotein, LDL: low-density lipoprotein, CI: confidence interval, N SNPs: number of single nucleotide polymorphisms.

## Discussion

We present the first large-scale GWAS of MD in an ancestrally diverse sample, including data from almost 1 million participants of African, East Asian and South Asian ancestry and Hispanic/Latinx samples. The largest previous report included 26,000 cases of African ancestry^19^. No large study so far has considered multiple ancestry groups simultaneously.

By aggregating data in ancestry-specific meta-analyses, we identified two novel loci. The association at 6q16 in the GWAS in samples of African ancestry requires further confirmation in future studies. However, the link with MD is biologically plausible. The lead variant was significantly associated with the expression of *MCHR2* specifically in brain cortex tissue. Melanin-concentrating hormone (MCH) is a neuropeptide that is expressed in the central and peripheral nervous systems. It acts as a neurotransmitter or neuromodulator in a broad array of neuronal functions directed toward the regulation of goal-directed behaviour, such as food intake, and general arousal^33^. MCH acts on two receptors, MCHR1 and MCHR2. Variants at *MCHR1* have been reported to be associated with bipolar disorder by a previous GWAS^34^. Moreover, the ASCC3 gene was observed as colocalised for both European and African ancestry-associated depression. It is involved in NF-kappa and AP1 transactivation^35^ and was reported in one of the largest TWASs of depression^19^ prioritised for cerebellum tissue. ASCC3 is thought to play a role for the circadian clock^36^ and has previously been associated with depression and anxiety comorbidity^37^.

The diversity in combination with large sample size enabled a comparison of the causal genetic architecture across ancestry groups. We assessed to what extent the 206 previously identified loci from large European ancestry discovery GWAS were transferable to other ancestry groups. Differences in allele frequencies, linkage disequilibrium between ancestry groups, and variable sample sizes impacted our power to observe associations for each group. We recently developed power-adjusted transferability ratios, an approach to account for all these factors by comparing observed transferable loci to what is expected for a study of a given ancestry and sample size^28^. The PAT ratios were about 30% for African, South Asian, and East Asian ancestry, remarkably similar and consistently low. We previously computed PAT ratios for several other traits and found variation between traits, but the estimates for MD were at the bottom^23,28^. With a PAT ratio of 64%, the transferability of MD loci discovered in European ancestry samples was much higher for the Hispanic/Latinx group. This finding may reflect that the Hispanic/Latinx group contained many participants with a high proportion of European ancestry^38,39^.

To better understand mechanisms underlying individual differences in vulnerability to development of MD, we need to bridge the gap from locus discovery to the identification of target genes. Our study achieved significant progress in this respect. Fine-mapping benefitted from the additional diverse samples, with median credible sets reduced from 65.6 to 35 in size and with 32 loci resolved to ≤10 putatively causal SNVs (11 loci from the European ancestry fine-mapping).

In the TWAS, expression of 354 genes was significantly associated with MD. Out of these, 205 gene associations were novel, and 89 were overlapping with results of the largest previously published MD TWAS^19^, which used S-MultiXcan for their analyses. Furthermore, 80 features were overlapping with hits from another, previously published, large MD TWAS with largely overlapping samples of European ancestry^32^, including *NEGR1, TRMT61A, BAG, TMEM106B, CKB, ESR2*. A number of these TWAS features (including *NEGR1, ESR2* and *TMEM106B*) were previously also fine-mapped (using FOCUS, a TWAS fine-mapping tool) and were highlighted as causal or putatively causal in previous post-TWAS analyses, strengthening the role of TWAS as an important tool to better understand the relationship between gene expression and MD.

Through TWAS and three other tools that incorporate the growing body of knowledge about functional annotations of the genome, we classified 43 genes as “high-confidence”. The definition admittedly remains arbitrary until the field establishes clear guidelines. Nevertheless, the high confidence list represents an evidence-based starting point for further follow-up. It provides confirmation for several genes that have repeatedly been highlighted as being near a GWAS associated variant and having high biological plausibility^6,17–19,32^: *NEGR1, DRD2, CELF4, LRFN5, TMEM161B*, and *TMEM106B*.

Furthermore, genes cadherin-9 *(CDH9)*, and protocadherins *(PCDHA1, PCDHA2, PCDHA3)* were classified as high-confidence. Cadherins are transmembrane proteins, mediating adhesion between cells and tissues in organisms^40^. In previous studies, cadherins have been linked with MD and with other disorders involving the brain, including late-onset Alzheimer’s disease (which often manifests as neuropsychiatric symptoms coupled with depression and anxiety)^41^. Howard and colleagues reported a putative association between cadherin-13 and depression^17^ and the involvement of the gene in ADHD, a neurodevelopmental disorder. Furthermore, two other studies investigating MD and other major mood disorders reported gene protocadherin-17 (protocadherins are the largest sub-family of cadherin-related molecules) as a novel susceptibility gene^42^. Finally, a study on MD by Xiao and colleagues also suggested protocadherins (*PCDH9*) as a putative, novel risk gene for MD, showing that *PCDH9* expression levels in the brain were reduced in MD patients compared to controls^43^. These previous findings are consistent with the results of our study, strengthening the evidence for the involvement of cadherins and protocadherins in the aetiology of MD.

Genes newly implicated in MD development in our study highlight novel pathways, pinpoint potential new drug targets, and suggest opportunities for drug repurposing. Gene *NDUFAF3* encodes mitochondrial complex I assembly protein which is the main target of the drug metformin^44^, the first-line drug for treatment of type 2 diabetes. Research in model organisms has provided a tentative link to a reduction in depression and anxiety^45^. Furthermore, a recent study, using more than 360,000 samples from the UK Biobank found associations between *NDUFAF3* and mood instability, suggesting that energy dysregulation may play an important role in the physiology of mood instability^46^.

Previous Mendelian Randomization (MR) studies conducted in populations of European ancestry suggested a causal relationship of higher BMI increasing the odds of depression^47–49^. To our knowledge, evidence of a reverse causal association (i.e., MD increases the odds of higher BMI) had not been previously reported^6^. We also observed that the genetic liability to MD was associated with higher triglyceride levels, lower HDL cholesterol and higher LDL cholesterol in subjects of European ancestry, which were not significant in the only previous MR study of smaller statistical power^50^. In other ancestry groups, no significant relationship between BMI and MD was observed. Moreover, our MR analyses showed an effect of reduced triglycerides on increasing odds of MD in participants of East Asian ancestry. Therefore, we provide further evidence for an opposite direction of effect for the relationship between MD and metabolic traits in European and East Asian ancestry groups^23^. Instead of generalising findings about depression risk factors across populations, further studies are needed to understand how genetic and environmental factors contribute to the complex relationships across diverse ancestry groups.

Our study has limitations. Whilst it provides an improved window into the diversity of MD genetics, sample size varied greatly by ancestry group. The smallest group were individuals of South Asian ancestry. To characterise MD in these populations, future studies prioritising primary data collection in these populations are needed. To contribute to this, we are currently recruiting MD patients and controls from Pakistan into the DIVERGE study^51,52^. However, a concerted global effort to increase diversity in genetics will be necessary to fully address the issue^25^. This also applies to the lack of other omics data and other functional databases to support downstream analyses for ancestrally diverse GWAS, such as large resources for transcriptomics or proteomics in relevant tissues^53,54^. This may have impacted on our TWAS results because the RNA sequencing data was predominantly from participants of European ancestry.

In this study, we assigned individuals into ancestry groups. Whilst this enabled important insights, e.g. about transferability of MD loci, such categorical assignments are imprecise and some participants with admixed ancestry may still get excluded. In future research, we aim to implement different analytic strategies that are fully inclusive.

Most of the individuals included in our study live in the United States or in the United Kingdom. Many identify with minority ethnic groups and this could impact on their risk factors and disease manifestations, making this a key research priority. Yet, it remains important to increase global diversity and study MD genetics across environments.

This study utilised data from several existing cohorts and bioresources to achieve large sample sizes. This necessitated using different outcome definitions, covering self-administered symptoms questionnaires, electronic healthcare records, as well as structured clinical interviews. The potential advantages and disadvantages of these approaches have been extensively discussed in previous studies^6,10^. It may be possible that differences in outcome definitions impacted on the low levels of transferability of existing genetic findings to different ancestry groups. However, it is unlikely to fully explain them. In a previous study based on strictly defined clinical major depressive disorder in China, the multi-ancestry genetic correlation with MD in the PGC-MDD was only 0.33^22^, suggesting that other factors related to ancestry or environment may impact on transferability. Moreover, in our analyses the PAT ratio was low for samples of East Asian, South Asian and African ancestry, but considerably higher for the Hispanic/Latinx group. Nevertheless, these were largely population-based studies where MD was defined based on short self-reported questionnaires.

In conclusion, in this first large-scale multi-ancestry GWAS of MD, we found a notable proportion of loci that are specific to certain ancestry groups. We identified novel, biologically plausible associations that were missed in European ancestry analyses and demonstrated that large diverse samples can be important for the identification of target genes and putative mechanisms. These findings suggest that for MD, a heterogeneous condition with highly complex aetiology, increasing ancestral as well as global diversity in genetic studies may be particularly important to ensure discovery of core genes and to inform about transferability of findings across ancestry groups.

## Methods

### Participating studies

We included data from 21 studies with ancestrally diverse participants. Details including study design, genotyping and imputation methods and quality control for these studies had been described by previous publications (**Study Descriptions**). All participants have provided informed consent. All studies obtained ethical approvals from local ethics review boards.

For each study, a principal component analysis was carried out based on the genetic similarity of individuals. Individuals who clustered around a reference group with confirmed ancestry were assigned to that specific group and included in the association analysis. Except for the Hispanic/Latinx group which was based on self-reported ethnicity, individuals with admixture between the pre-defined ancestry reference groups were excluded.

We also included previously published studies of MD using data from ancestrally European participants, including PGC-MDD (N cases = 246,363, N controls = 561,190)^17^ and the AGDS (N cases = 12,123, N controls = 12,684)^18^ to conduct a multi-ancestry meta-analysis of MD). The total sample size of the multi-ancestry meta-analysis was 1,820,927 (N cases = 342,883; N_*effhalf*_ = 485,107). 70.1% of participants (effective sample size) were of European ancestry and 8.2% East Asian, 11.8% African and 1.5% South Asian ancestry and 7.9% Hispanic/Latinx.

### Study level genetic association analyses

We had access to individual level data for Army-STARRS, UKB, WHI, and IHS, GERA, JHS, Drakenstein Child Health Study, and the DNHS study. Data access was granted via our collaborators, the UK Biobank under application ID 51119 and dbGaP under project ID 18933. A range of measures were used to define depression, including structured clinical interviews, medical care records, symptom questionnaires and self-reported surveys (**Supplementary Table 12**). SNV-level associations with depression were assessed through logistic regressions using PLINK. The additive per-allele model was employed. Age, sex, principal components and other relevant study-level covariates were included as covariates. Where available, genotypes on chromosome X were coded 0 or 2 in male participants and 0, 1 and 2 in female participants. Summary statistics were received from our collaborators for all other studies. Additive-effect logistic regressions were conducted by the 23andMe Inc, Taiwan-MDD study, MVP, BBJ, Rabinowitz, MAMHS, PrOMIS and BioVU. Age, sex, principal components and other relevant study-level covariates were included as covariates.

Mixed-effect models were used in the association analysis for CKB, BioME, Genes & Health with SAIGE (version 0.36.1)^55^. The CONVERGE study initially conducted mixed-effect model GWA tests with bolt-LMM, followed by PLINK logistic regressions to retrieve logORs. For the CONVERGE study, the logORs and standard errors from PLINK were used in our meta-analysis. The HCHS/SOL implemented mixed-effect model GWA tests to adjust for population structure and relatedness with depression as binary outcome^20^ and was run using GENESIS^56^. The summary statistics from GENESIS were converted into logOR and SE before meta-analysis. First, the score and its variance were transformed into beta and se by beta = score/variance and se = sqrt(variance)/variance. Afterwards, beta and se were converted into approximate logOR and se using beta = beta/(pi*(1-pi)) and se = se/(pi*(1-pi)), where pi is the proportion of cases in analysis^57^.

We restricted the downstream analysis to variants with imputation accuracy info score of 0.7 or higher and effective allele count (2*maf*(1-maf)*N*R^2^) of 50 or higher. For study of small sample size, we instead required a minor allele frequency of no less than 0.05. The alleles for indels were re-coded as “I” for the longer allele and “D” for the shorter one. Indels of different patterns at the same position were removed.

### Meta-analyses

We first implemented inverse-variance weighted fixed-effect meta-analyses for GWAS from each ancestry/ethnic group (i.e., African ancestry, East Asian ancestry, South Asian ancestry and Hispanic/Latinx) using METAL^58^. The genomic inflation factor λ was calculated for each study and meta-analysis. Given the dependence of this estimate on sample size, we also calculated λ_1000_ ^59^ as λ_1000_ = 1 + (λ - 1) * (1 / Ncase + 1 / Ncontrol) * 500 ^60^. The LDSC intercept was also calculated with an ancestry matched LD reference panel from the PAN UKB reference panel^61^ for each meta-analysis^62^. For meta-analyses with residual inflation (λ > 1.1), test statistics for variants were adjusted by LDSC intercept. Following the meta-analyses by METAL, variants present in less than 2 studies were filtered out.

In order to facilitate multi-ancestry analyses, a multiple ancestry LD reference panel was constructed by randomly drawing 10k participants from the UK Biobank. The proportions of individuals from each of the four major ancestries (European, African, East Asian and South Asian) in the multiple ancestry LD reference panel was matched to the proportions of ancestries in our multi-ancestry MD GWAS on the effective sample size scale. Hispanic/Latinx descent participants were omitted from the multiple ancestry LD reference due to their complex structure and low representation in the UK Biobank data. As a result, there were 7,658 participants of European (76.58%), 1,285 of African (12.85%), 891 of East Asian (8.91%) and 166 of South Asian (1.66%) ancestry in the multi-ancestry LD reference panel.

We combined data from 71 cohorts with diverse ancestry using an inverse-variance weighted fixed-effects meta-analysis in METAL^58^. λ and λ_1000_ were calculated, which were 1.687 and 1.001, respectively. The LDSC intercept was also calculated with the multi-ancestry LD reference panel, which was 1.019 (SE = 0.011). We adjusted the test statistics from the multi-ancestry meta-analysis using the LDSC intercept of 1.019. Only variants present in at least two studies were retained for further analysis, yielding a total of 22,941,580 variants. We also calculated the number of cases and the total number of samples for each variant based on the crude sample size and availability of each study.

We used a significance threshold of 5×10^−8^. To identify independent association signals, the GCTA forward selection and backward elimination process (command ‘cojo-slt’) were applied using the summary statistics from the multi-ancestry meta-analysis, with the aforementioned multi ancestry LD reference panel^63,64^. It is possible that the algorithm identifies false positive secondary signals if the LD in the reference set does not match the actual LD in the GWAS data well. Therefore, for each independent signal defined by the GCTA algorithm, locus zoom plots were generated for the 250kb upstream and downstream region. We then inspected each of these plots manually and removed any secondary signals from our list where there was unclear LD separation, i.e. some of the variants close to the secondary hit were in LD with the lead variant.

Loci were defined by the flanking genomic interval mapping 250kb upstream and downstream of each lead SNV. Where lead SNVs were separated by less than 500kb, the corresponding loci were aggregated as a single locus with multiple independent signals. The lead SNV for each locus was then selected as the SNV with minimum association P value.

### Fine-mapping

We fine-mapped all loci with statistically significant associations from the multi-ancestry GWAS using a statistical fine-mapping method for multi-ancestry samples^29^. In short, this method is an extension of a Bayesian fine-mapping approach^29,65^ that utilises estimates of the heterogeneity across ancestry groups such that variants with different effect estimates across populations have a smaller prior probability to be the causal variant.

For each lead variant, we first extracted all nearby variants with *r*^*2*^ > 0.1 as determined by the multi-ancestry LD reference. The multi-ancestry prior for each variant to be causal was calculated from a fixed effects meta-analysis combining the summary statistics from ancestry-specific meta-analysis for each of the five major ancestry groups. *I*^*2*^ statistics were calculated to estimate the heterogeneity of the effect estimates across ancestry groups. The posterior probability for a variant to be included in the credible set was proportional to its chi-square test statistic and the prior. The 99% credible set for each lead variant was determined by ranking all SNVs (within *r*^*2*^ > 0.1 of the lead variant) according to their posterior probabilities and then including ranked SNVs until their cumulative posterior probabilities reached or exceeded 0.99.

As a comparison, we also conducted a Bayesian fine-mapping analysis based on the summary statistics of the European-ancestry meta-analysis. The same list of independent lead SNVs from the multi-ancestry meta-analysis were used for this fine-mapping in the European ancestry data. All nearby SNVs with *r*^*2*^ > 0.1 as determined by the 1000 Genomes European LD reference panel were included in the fine-mapping. The posterior probability was calculated in a similar way, but without the multi-ancestry prior. Similar to the multi-ancestry fine-mapping, all SNVs were ranked and 99% of the credible sets were derived accordingly.

Since our fine-mapping was based on meta-analysis summary statistics, heterogeneity of individual studies (e.g., due to differences in genotyping array) can influence the fine-mapping calibration and recall. We used a novel summary statistics-based QC method proposed by Kanai and colleagues (SLALOM) to dissect outliers in association statistics for each fine-mapped locus^66^. This method calculates test statistics (DENTIST-S) from Z-scores of test variants and the lead variant (the variant of the lowest P value in each locus), and the LD r between test variants and the lead variant in the locus^67^. Among the 155 fine-mapped loci in our study, there were 134 loci with the largest variant posterior inclusion probability of greater than 0.1. For these 134 loci, *r* values were calculated for all variants within the 1MB region of the lead variant for each locus based on our multi-ancestry LD reference from the UK Biobank data. In line with the criteria used by Kanai and colleagues, variants with DENTIST-S *P* value smaller than 1×10^−4^ and *r*^*2*^ with the lead variant greater than 0.6 were defined as outliers. Fine-mapped loci were classified as robust if there were no outlying variants.

### Colocalisation analysis

We performed colocalization between genetic associations with MD and gene expression in brain and blood tissues from samples of European and African ancestry and Hispanic/Latinx participants using coloc R package^68^. To select genes for testing, we mapped SNVs within a 3MB window at 2q24.2 and 6q16.2 using Variant Effect Predictor^30^, resulting in eight and four genes, respectively. Loci with posterior probability >90% either for, both traits are associated and share two different but linked variants (H3 hypothesis), or a single causal variant (H4 hypothesis) were considered as colocalized. The European and African ancestry summary statistics for MD were tested against multi-ancestry brain eQTLs from European and African American samples^27^. For Hispanic/Latinx ancestry, we tested gene and protein expression of blood tissue from Multi-Ethnic Study of Atherosclerosis and Trans-omics for Precision Medicine^69^. For African ancestry, we tested gene expression of blood from GENOA study^70^ and proteome expression of blood^71^. For European ancestry, we tested gene expression of blood from eQTLgen^72^, and proteome expression from blood^71^.

### Assessment of transferability of established loci

We assessed whether published MD-associated loci display evidence of association in the East Asian, South Asian, African ancestry and Hispanic/Latinx samples. Pooling the independent genome-wide significant SNVs from two large GWAS of MD in samples of European ancestry yielded 195 loci^17–19^. The ancestrally diverse groups included in this study had smaller numbers of participants than the European ancestry discovery studies. Also, a given variant may be less frequent in another ancestry group. Therefore, individual lead variants may not display evidence of association because of lack of power. Moreover, in the discovery study the lead variant is either the causal variant or is strongly correlated with it. However, differences in LD mean that the lead variant may not be correlated in another ancestry group and may therefore not display evidence of association. Our assessment of transferability was therefore based on power-adjusted transferability (PAT) ratios that aggregate information across loci and account for all three factors, sample size, MAF and differences in LD^28^.

First, credible sets for each locus were generated. They consisted of lead variant plus all correlated SNVs (r^2^ >= 0.8) within a 50kb window of the lead variant (based on ancestry matched LD reference panels from the 1000 Genomes data) and with *P* value < 100 × *P*_lead_. A signal was defined as being ‘transferable’ to another ancestry group if at least one variant from the credible set was associated at two-sided *P* < 10^(log(0.05,base=10)-P_f_ × (N-1)) with MD and consistent direction of effect between the discovery and test study. N is the number of SNVs in the credible set for each locus, and P_f_ is a penalization factor we derived from empirical estimations. The effective number of independent SNVs was often higher in other ancestry groups due to differences in LD, leading to higher multiple testing burden and higher likelihood of identifying SNVs with a low p-value, by chance alone. This inflates the test statistics and was adjusted for by the penalization factor (P_f_). In order to derive the P_f_ for each ancestry group, we used the summary statistics from a previous GWAS on breast cancer^73^, in which phenotypes were believed to be uncorrelated with MD. A total of 441 breast cancer significant SNVs were taken from their paper and linear regressions were conducted for the *P* values of these SNVs in each of our ancestrally diverse summary statistics for MD on the number of SNVs in credible sets. The coefficient estimates (slope from regressions) were treated as P_f_ for each ancestry. As a result, P_f_ were 0.008341, 0.007378, 0.006847 and 0.003147 for samples of African, East Asian, South Asian ancestry and Hispanic/Latinx, respectively.

In the next step, the statistical power to detect an association of a given locus was calculated assuming an additive effect at a type I error rate of 0.05, with effect estimates from the discovery study, and allele frequency and sample size from each of the target data sets from diverse ancestry/ethnic groups. The power estimates were summed up across published loci to give an estimate of the total number of loci expected to be significantly associated. This is the expected number if all loci are transferable and accounts for the statistical power for replication. We calculated the PAT ratio by dividing the observed number of loci by the expected number. In addition, loci were defined as ‘non-transferable’ if they had sufficient power for identifying an association but did not display evidence of association, ie., if they contained at least one variant in the credible set with > 80% power, while none of the variants in the credible set had *P* < 0.05 and no variant within 50kb of locus had *P* < 1×10^−3^ in the target data set.

For comparison, we also conducted a transferability assessment for a European ancestry look-up study. The 102 significant loci reported by Howard and colleagues^17^ were evaluated for their transferability in the AGDS study using the aforementioned method.

### Gene Annotation

The summary statistic from the multi-ancestry meta-analysis was first annotated with FUMA^74^. Both positional mapping and eQTL mapping results were extracted from FUMA. The 1000 Genomes Europeans were employed as the LD reference panel for FUMA gene annotation. Data sets for brain tissue available in FUMA were employed for eQTL gene annotation.

Gene-based association analyses were implemented using Multi-marker Analysis of GenoMic Annotation (MAGMA, v1.08)^75^ and Hi-C coupled MAGMA (H-MAGMA)^76^. The aforementioned multiple ancestry LD reference panel from the UK Biobank was used as the LD reference panel. H-MAGMA assigns non-coding SNVs to their cognate genes based on long range interactions in disease-relevant tissues measured by Hi-C ^76^. We used the adult brain Hi-C annotation file.

### Transcriptome-wide association analysis and drug mapping

To perform a transcriptome-wide association study (TWAS), the FUSION software was used^77^. SNP-weights were downloaded from the FUSION website^78^ and were derived from multiple external studies, including:

1. SNP-weights from all available brain tissues, adrenal gland, pituitary gland, thyroid gland and whole blood^32^ from GTEx v8, (based on significantly heritable genes and “All Samples” in GTEx v8, which also includes African American and Asian individuals^79^.
2. SNP-weights from the CommonMind Consortium (CMC), which includes samples from the brain dorsolateral prefrontal cortex.
3. and SNP-weights from the Young Finns study (YFS) and
4. Netherlands Twin Register (NTR) which provides SNP-weights from blood tissues (whole blood and peripheral blood, respectively).

We used the multi-ancestry LD reference panel described above. Variants present in the 1000 Genomes European population reference panel were retained. A separate TWAS was also performed using a LD reference panel based on the 1000 Genomes Project’s samples of European ancestry, as a sensitivity analysis.

The transcriptome-wide significance threshold for the TWAS associations in this study was *P*<1.37×10^−6^. This threshold has previously been derived using a permutation-based procedure, which estimates a significance threshold based on the number of features tested^32^.

Results were compared with previous TWASs in MD, including the two largest MD TWASs to date^6^,^80^,^81^,^32^,^19^. These studies generally used smaller sets of SNP-weights (except the study by Dall’Aglio and colleagues, which used similar SNP-weights as the current study, but with SNP-weights derived from the previous GTEx release, v7). A TWAS z-score plot was generated using a TWAS-plotter function^82^.

To assess the relevance of novel genes to drug discovery, genes were searched in three large drug databases: in Genecards.org, DrugBank and ChEMBL^83^,^84^,^85^. In Table 1, a selection of drugs (the ones reported in multiple publications) likely targeting our high confidence prioritised gene sets are shown for each gene.

### Mendelian Randomisation

We performed a bi-directional two-sample MR analysis using the TwoSampleMR R package^86^ to test possible causal effects between MD and six cardiometabolic traits. For individuals of European ancestry, the UK Biobank was used to select instruments for BMI, fasting glucose (FG), high-density lipoprotein (HDL), low-density lipoprotein (LDL), systolic blood pressure (SBP), and triglycerides (TG). SBP summary data were obtained from the UK Biobank for individuals of African and South Asian ancestry and Hispanic/Latinx participants. For samples of African, East Asian and South Asian ancestry and the Hispanic/Latinx group, a meta-analysis was performed using METAL^58^ with inverse variance weighting using the UK Biobank and the following consortia: GIANT^87^ for BMI, MAGIC^88^ for FG, GLGC^89^ for HDL, LDL, and TG, and BBJ^89,90^ for SBP in samples of East Asian ancestry. To avoid sample overlap, the datasets used to define instrumental variables for the cardiometabolic traits were excluded from the MD genome-wide association statistics used for the MR analyses conducted with respect to each ancestry group.

Genome-wide significance (*P* = 5×10^−8^) was used as the threshold to select instrumental variables for the exposures. However, if less than 10 variants were available, a suggestive threshold (*P* = 5×10^−6^) was used to select instrumental variables **(Supplementary Table 11)**. We used five different MR methods: inverse variance weighted (IVW), MR-Egger, weighted median, simple mode, and weighted mode^86^. The IVW estimates were reported as the main results due to their higher statistical power^91^ while the other tests were used to assess the consistency of the estimates across different methods. MR-Egger regression intercept and MR heterogeneity tests were conducted as additional sensitivity analyses. In case of significant heterogeneity, the MR-PRESSO (Pleiotropy RESidual Sum and Outlier) global test was used to remove genetic variants based on their contribution to heterogeneity^92^).

## Acknowledgements

We are grateful to all the participants who took part in the studies and would also like to acknowledge the investigators involved in the participating studies. We would like to thank all members of the ULC HumGen lab (https://www.uclhumgen.com/), who gave critical support and suggestions. We would like to thank Albert Henry, MD, University College London, for discussions and suggestions in key analytic techniques. This work was conducted on the University College London Computer Science cluster, we would like to thank the cluster team for the support provided. **China Kadoorie Biobank**’s most important acknowledgement is to the participants in the study. The investigators also acknowledge the invaluable contributions of the members of the survey teams in each of the 10 regional centres, and of the project development and management teams based at Beijing, Oxford, and the 10 regional centres. China’s National Health Insurance provides electronic linkage to all hospital treatments.. **Mexican Adolescent Mental Health Survey cohort** thank Corina Benjet and Enrique Méndez for their contributions to the Mexican Adolescent Mental Health Survey cohort’s data acquisition and curation, respectively. **Intern Health Study** thank the training physicians for taking part in Intern Health Study. This project was funded by the National Institute of Mental Health (grant no. R01MH101459). **MVP** thank the veterans who participate in the Million Veteran Program. From the Yale Department of Psychiatry, Division of Human Genetics, we would like to thank and acknowledge the efforts of A. M. Lacobelle, C. Robinson and C. Tyrell. Funding: this work was supported by funding from the Veterans Affairs Office of Research and Development Million Veteran Program grant CX001849-01 (MVP025) and VA Cooperative Studies Program CSP575B. D.F.L. was supported by an NARSAD Young Investigator Grant from the Brain & Behavior Research Foundation. **The BioVU study** used for the analyses described were obtained from Vanderbilt University Medical Center’s BioVU which is supported by numerous sources: institutional funding, private agencies, and federal grants. These include the NIH funded Shared Instrumentation Grant S10RR025141; and CTSA grants UL1TR002243, UL1TR000445, and UL1RR024975. Genomic data are also supported by investigator-led projects that include U01HG004798, R01NS032830, RC2GM092618, P50GM115305, U01HG006378, U19HL065962, R01HD074711; and additional funding sources listed at https://victr.vumc.org/biovu-funding/. **CONVERGE a**uthors are part of the CONVERGE consortium (China, Oxford and Virginia Commonwealth University Experimental Research on Genetic Epidemiology) and gratefully acknowledge the support of all partners in hospitals across China. Special thanks to all the CONVERGE collaborators and patients who made this work possible. CONVERGE Consortium: Na Cai, Tim B. Bigdeli, Warren Kretzschmar, Yihan Li, Jieqin Liang, Li Song, Jingchu Hu, Qibin Li, Wei Jin, Zhenfei Hu, Guangbiao Wang, Linmao Wang, Puyi Qian, Yuan Liu, Tao Jiang, Yao Lu, Xiuqing Zhang, Ye Yin, Yingrui Li, Xun Xu, Jingfang Gao, Mark Reimers, Todd Webb, Brien Riley, Silviu Bacanu, Roseann E. Peterson, Yiping Chen, Hui Zhong, Zhengrong Liu, Gang Wang, Jing Sun, Hong Sang, Guoqing Jiang, Xiaoyan Zhou, Yi Li, Yi Li, Wei Zhang, Xueyi Wang, Xiang Fang, Runde Pan, Guodong Miao, Qiwen Zhang, Jian Hu, Fengyu Yu, Bo Du, Wenhua Sang, Keqing Li, Guibing Chen, Min Cai, Lijun Yang, Donglin Yang, Baowei Ha, Xiaohong Hong, Hong Deng, Gongying Li, Kan Li, Yan Song, Shugui Gao, Jinbei Zhang, Zhaoyu Gan, Huaqing Meng, Jiyang Pan, Chengge Gao, Kerang Zhang, Ning Sun, Youhui Li, Qihui Niu, Yutang Zhang, Tieqiao Liu, Chunmei Hu, Zhen Zhang, Luxian Lv, Jicheng Dong, Xiaoping Wang, Ming Tao, Xumei Wang, Jing Xia, Han Rong, Qiang He, Tiebang Liu, Guoping Huang, Qiyi Mei, Zhenming Shen, Ying Liu, Jianhua Shen, Tian Tian, Xiaojuan Liu, Wenyuan Wu, Danhua Gu, Guangyi Fu, Jianguo Shi, Yunchun Chen, Xiangchao Gan, Lanfen Liu, Lina Wang, Fuzhong Yang, Enzhao Cong, Jonathan Marchini, Huanming Yang, Jian Wang, Shenxun Shi, Richard Mott, Qi Xu, Jun Wang, Kenneth S. Kendler & Jonathan Flint. The **Australian Genetics of Depression Study** are indebted to all of the participants for giving their time to contribute to this study. We wish to thank all the people who helped in the conception, implementation, media campaign and data cleaning. We thank Richard Parker, Simone Cross and Lenore Sullivan for their valuable work coordinating all the administrative and operational aspects of the AGDS project. **The 23andMe Study** would like to thank the research participants and employees of 23andMe for making this work possible. The following members of the 23andMe Research Team contributed to this study: Stella Aslibekyan, Adam Auton, Elizabeth Babalola, Robert K. Bell, Jessica Bielenberg, Katarzyna Bryc, Emily Bullis, Daniella Coker, Gabriel Cuellar Partida, Devika Dhamija, Sayantan Das, Sarah L. Elson, Nicholas Eriksson, Teresa Filshtein, Alison Fitch, Kipper Fletez-Brant, Pierre Fontanillas, Will Freyman, Julie M. Granka, Karl Heilbron, Alejandro Hernandez, Barry Hicks, David A. Hinds, Ethan M. Jewett, Katelyn Kukar, Alan Kwong, Keng-Han Lin, Bianca A. Llamas, Maya Lowe, Jey C. McCreight, Matthew H. McIntyre, Steven J. Micheletti, Meghan E. Moreno, Priyanka Nandakumar, Dominique T. Nguyen, Elizabeth S. Noblin, Jared O’Connell, Aaron A. Petrakovitz, G. David Poznik, Alexandra Reynoso, Morgan Schumacher, Anjali J. Shastri, Janie F. Shelton, Jingchunzi Shi, Suyash Shringarpure, Qiaojuan Jane Su, Susana A. Tat, Christophe Toukam Tchakouté, Vinh Tran, Joyce Y. Tung, Xin Wang, Wei Wang, Catherine H. Weldon, Peter Wilton, Corinna D. Wong. **Genes & Health** We thank Social Action for Health, Centre of The Cell, members of our Community Advisory Group, and staff who have recruited and collected data from volunteers. We thank the NIHR National Biosample Centre (UK Biocentre), the Social Genetic & Developmental Psychiatry Centre (King’s College London), Wellcome Sanger Institute, and Broad Institute for sample processing, genotyping, sequencing and variant annotation. We thank: Barts Health NHS Trust, NHS Clinical Commissioning Groups (City and Hackney, Waltham Forest, Tower Hamlets, Newham, Redbridge, Havering, Barking and Dagenham), East London NHS Foundation Trust, Bradford Teaching Hospitals NHS Foundation Trust, Public Health England (especially David Wyllie), Discovery Data Service/Endeavour Health Charitable Trust (especially David Stables) - for GDPR-compliant data sharing backed by individual written informed consent. Most of all we thank all of the volunteers participating in Genes & Health.

## Funding

This study is part of a project that has received funding from Wellcome (212360/ Z/18/Z) and from the European Research Council (ERC) under the European Union’s Horizon 2020 research and innovation program (Grant agreement No. 948561). Computing was supported by the BBSRC Biotechnology and Biological Sciences Research Council (BB/R01356X/1). D.K. is supported by the MSCA Individual Fellowship, European Commission (101028810). G.N. is supported by Biotechnology and Biological Sciences Research Council (BBSRC) grant number BB/M009513/1. R.P. is supported by Grants from the National Institutes of Health (R33 DA047527; R21 DC018098) and One Mind Rising Star Award. G.P is supported by the Yale Biological SciencesTraining Program (T32 MH014276). P.H.K is supported by The Ministry of Science and Technology Project (MOST 108-2314-B-002-136-MY3), the National Health Research Institutes Project (NHRI-EX106-10627NI), and The National Taiwan University Career Development Project (109L7860). The AGDS was primarily funded by the National Health and Medical Research Council (NHMRC) of Australia grant 1086683. This work was further supported by NHMRC grants 1145645, 1078901, 1113400 and 1087889 and NIMH. The QSkin study was funded by the NHMRC (grant numbers 1073898, 1058522 and 1123248). N.G.M. is supported through NHMRC investigator grant 1172990. N.R.W. is supported through NHMRC investigator grant 1173790. W.E. is supported by NIDA grant R01DA009897. CONVERGE was funded by the Wellcome Trust (WT090532/Z/09/Z, WT083573/ Z/07/Z, WT089269/Z/09/Z) and by NIH grant MH100549. K.S.K. was supported by NIMH R01MH125938 and R21MH126358. R.E.P. was supported by NIMH R01MH125938, R21MH126358, and The Brain & Behavior Research Foundation NARSAD grant 28632 P&S Fund. L.K.D. is supported by funding R01 MH118223. R.J.U., R.C.K., M.B.S. is supported by the US Department of Defense. J.G. and the MVP study was supported by funding from the Veterans Affairs Office of Research and Development Million Veteran Program grant CX001849-01 (MVP025) and VA Cooperative Studies Program CSP575B. S.S. is supported by R01MH101459. M.U., D.W., and A.A. are supported by 2R01MD011728. C.S.C-F. is supported by Cohen Veteran Bioscience, Mexico’s National Institute of Psychiatry, Mexico’s National Council of Science & Technology (CONACyT). The Hispanic Community Health Study/Study of Latinos is a collaborative study supported by contracts from the National Heart, Lung, and Blood Institute (NHLBI) to the University of North Carolina (HHSN268201300001I / N01-HC-65233), University of Miami (HHSN268201300004I / N01-HC-65234), Albert Einstein College of Medicine (HHSN268201300002I / N01-HC-65235), University of Illinois at Chicago (HHSN268201300003I / N01-HC-65236 Northwestern Univ), and San Diego State University (HHSN268201300005I / N01-HC-65237). The following Institutes/Centers/Offices have contributed to the HCHS/SOL through a transfer of funds to the NHLBI: National Institute on Minority Health and Health Disparities, National Institute on Deafness and Other Communication Disorders, National Institute of Dental and Craniofacial Research, National Institute of Diabetes and Digestive and Kidney Diseases, National Institute of Neurological Disorders and Stroke, NIH Institution-Office of Dietary Supplements. The Genetic Analysis Center at the University of Washington was supported by NHLBI and NIDCR contracts (HHSN268201300005C AM03 and MOD03). The Hispanic Community Health Study/Study of Latinos also received support from the National Institutes of Health Award (R01MH113930). B.S.M. and W.E. is supported by the NIDA. J.A.R. and G.R.U. are supported by NIDA, NIA, VA, University of Maryland, Maryland VA Healthcare System, Baltimore Research and Education Foundation. The China Kadoorie Biobank baseline survey and the first re-survey were supported by the Kadoorie Charitable Foundation in Hong Kong. Long-term follow-up was supported by the Wellcome Trust (212946/Z/18/Z, 202922/Z/16/Z, 104085/Z/14/Z, 088158/Z/09/Z), the National Key Research and Development Program of China (2016YFC0900500, 2016YFC0900501, 2016YFC0900504, 2016YFC1303904), and the National Natural Science Foundation of China (91843302). DNA extraction and genotyping was funded by GlaxoSmithKline, and the UK Medical Research Council (MC-PC-13049, MC-PC-14135). The China Kadoorie Biobank is supported by core funding from the UK Medical Research Council (MC_UU_00017/1,MC_UU_12026/2, MC_U137686851), Cancer Research UK (C16077/A29186; C500/A16896), and the British Heart Foundation (CH/1996001/9454) to the Clinical Trial Service Unit and Epidemiological Studies Unit and to the MRC Population Health Research Unit at Oxford University. Genes & Health has recently been core-funded by Wellcome (WT102627, WT210561), the Medical Research Council (UK) (M009017), Higher Education Funding Council for England Catalyst, Barts Charity (845/1796), Health Data Research UK (for London substantive site), and research delivery support from the NHS National Institute for Health Research Clinical Research Network (North Thames). Genes & Health has recently been funded by Alnylam Pharmaceuticals, Genomics PLC; and a Life Sciences Industry Consortium of Bristol-Myers Squibb Company, GlaxoSmithKline Research and Development Limited, Maze Therapeutics Inc, Merck Sharp & Dohme LLC, Novo Nordisk A/S, Pfizer Inc, Takeda Development Centre Americas Inc.

## Author Contributions

K.K. conceived this project and supervised the work. X.M. and O.G. carried out the study-level GWAS analysis. Y.F. carried out meta-analysis for studies of African ancestry. X.M. carried out all meta-analysis, loci and independent SNV identification, transferability analysis, fine mapping, and annotation including FUMA, MAGMA and Hi-C MAGMA. G.N. carried out the TWAS analysis and conducted the drug target look-up. D.K. and R.P. carried out the Mendelian Randomization analysis. G.P. and R.P. carried out the QTL and colocalization analysis. A.M. provided support for analyses and result interpretation. X.M., G.N., and K.K. wrote and critically revised the manuscript. M.V. contributed to drafting and editing of the manuscript. MVP: J.G and M.B.S. are the principal investigators, J.M.G. and D.L. were involved in preparing data. China Kadoorie Biobank: Z.C. and L.L. are the principal investigators; R.G.W. is the genomics lead; R.G.W., I.Y.M. were involved in data collection; R.G.W. and K.L. carried out quality control and genome-wide association analysis; K.L. carried out the genotype imputation. Genes & Health: D.A.v.H. is the principal investigator, B.T., S.F., N.B., H.M., V.K.C and Q.Q.H. were involved in data collection and analysis. The GREAT study cohort: P.H.K is the principal investigator, H.C.C, S.J.T. and Y.L.L. were involved in data collection and analysis. AGDS: N.G.M, N.R.W. and E.M.B. are the principal investigators, B.L.M. carried out data analysis. CONVERGE: K.S.K. is the principal investigator, R.E.P. and N.C. were involved in data collection and analysis. BioVU: L.K.D. is the principal investigator, K.V.A were involved in data collection and analysis. Army STARRS: R.J.U. and M.B.S. are the principal investigators, and R.C.K. was involved in data collection and preparation. IHS: S.S. is the principal investigator, Y.F., L.S. and M.B. were involved in data collection and preparation. BioMe: R.L. is the principal investigator and M.P. was involved in data collection and analysis. 23andMe: Y.J. and C.T. were involved in data collection and analysis. Drakenstein Child Health Study: D.J.S and H.J.Z. are the principal investigator, M.L.C. and N.K. were involved in data collection and preparation. Detroit Neighborhood Health Study: M.U. is the principal investigator, A.W. D.W. and A.A. were involved in data collection and preparation. Mexican Adolescent Mental Health Survey Cohort: C.S.C-F. is the principal investigator, G.A. M-L., M.E.R. and A.I.C. were involved in data collection and analysis. Hispanic Community Health Study/Study of Latinos: E.C.D. and S.W-S. are the principal investigators, T.S. was involved in data collection and analysis. PIRC 1st generation trial: W.E. is the principal investigator, J.A.R., B.S.M., and G.R.U. were involved in data collection and analysis. Biobank Japan: Y.O. is the principal investigator, M.K. and S.S. were involved in data collection and analysis. PGC MDD: S.A., J.R.I.C., and S.R. contributed to the PGC MDD study of European ancestry. All authors read and critically revised the manuscript and made significant intellectual contributions to the study. All authors approved the manuscript.

## Conflict of Interest

None

## Notes

### Competing Interest Statement

The authors have declared no competing interest.

