## Supplementary Figures for "Multi-ancestry GWAS of major depression aids locus discovery, fine-mapping, gene prioritisation, and causal inference"

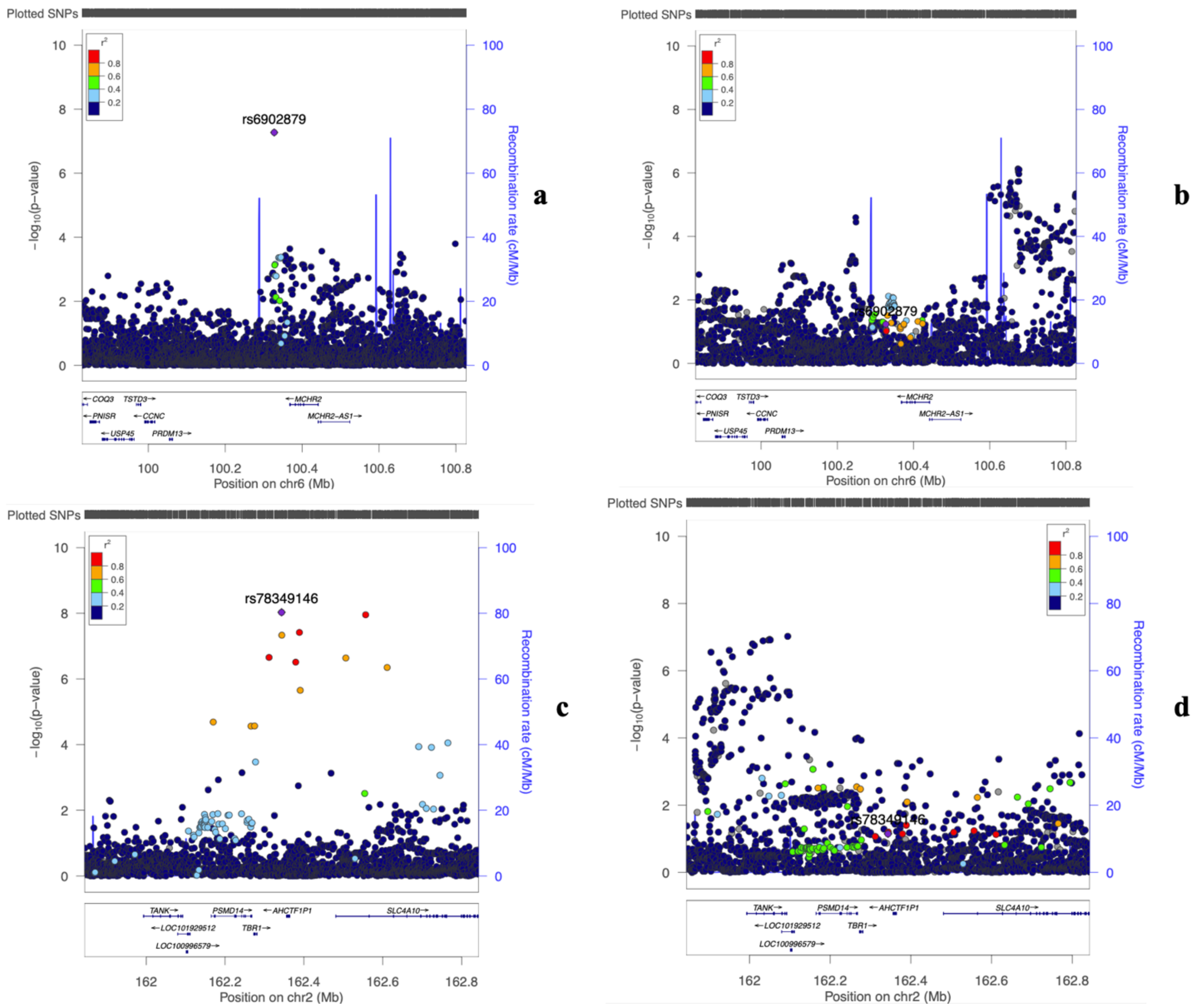


### Supplementary Figure 1. Regional association plots for the associations with major depression for two loci from ancestry-specific GWAS.

The y-axes show the -log10P values of the association between each SNV and the outcome. The x-axes show the chromosomal position (GRCh37). A. Genetic associations for rs6902879 and variants within 500kb region in African ancestry meta-analysis; B. Genetic associations for rs6902879 and variants within 500kb region in European ancestry meta-analysis; C. Genetic associations for rs78349146 and variants within 500kb region in meta-analysis of Hispanic/Latinx individuals; D. Genetic associations for rs78349146 and variants within 400kb region in European ancestry meta-analysis.

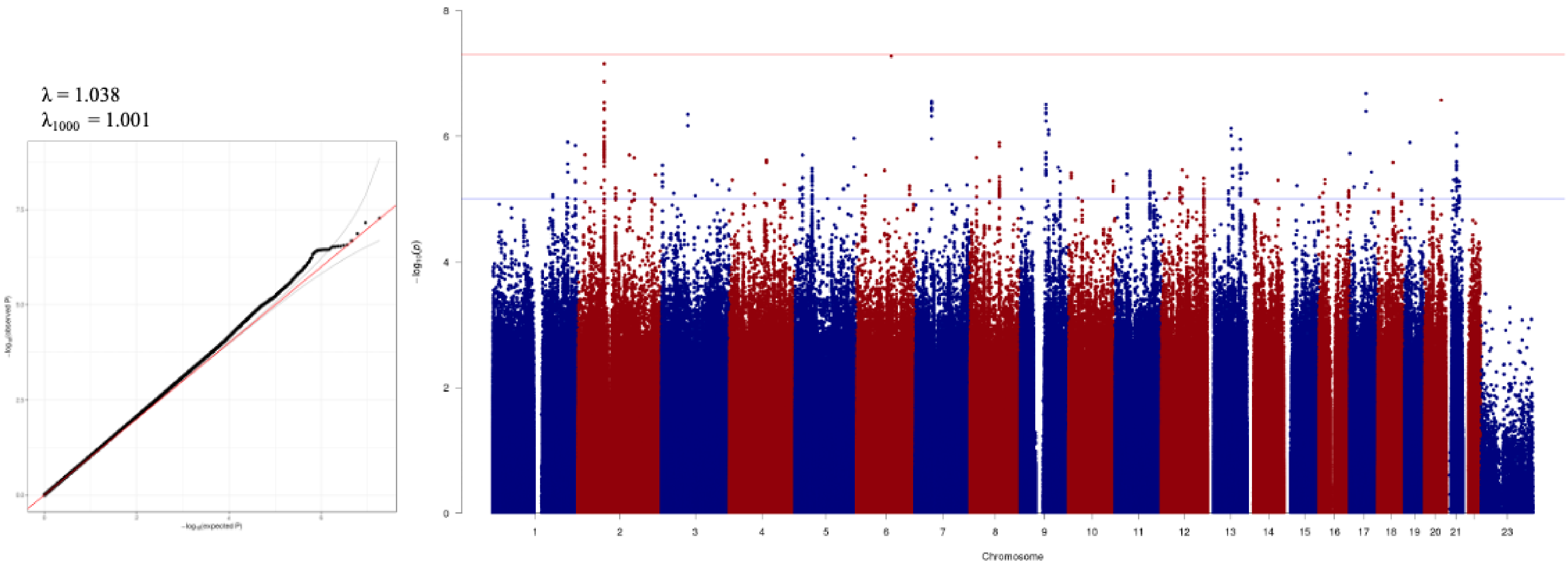


Supplementary Figure 2. Manhattan plot and QQ plot with depression in individuals of African ancestry.

The y-axes show the −log10*P* values of the association between each single-nucleotide variant and the outcome. The x-axes show the chromosomal position (GRCh37). The red line represents the genome-wide significance threshold of 5 x $10^{-8}$ and the blue line, $10^{-5}$.

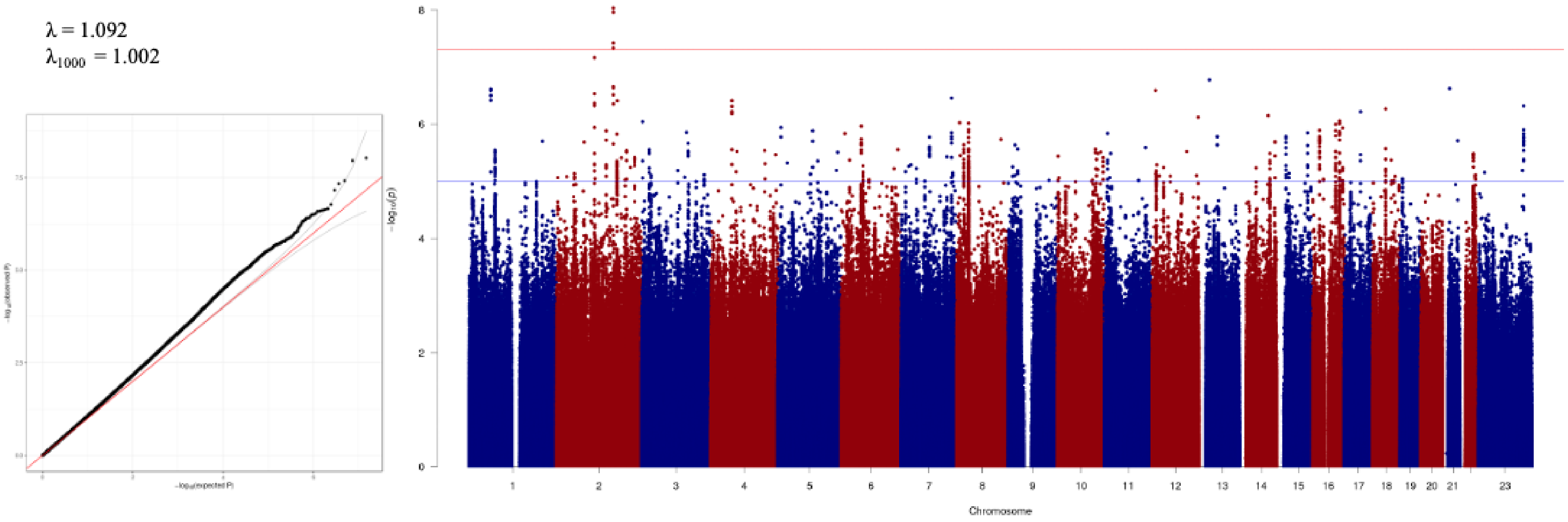


Supplementary Figure 3. Manhattan plot and QQ plot with depression in individuals of Latinx/Hispanic ancestry.

The y-axes show the −log10*P* values of the association between each single-nucleotide variant and the outcome. The x-axes show the chromosomal position (GRCh37). The red line represents the genome-wide significance threshold of $5 \times 10^{-8}$ and the blue line, $10^{-5}$. Association *P* values have been adjusted by the LDSC intercept of 1.0508.

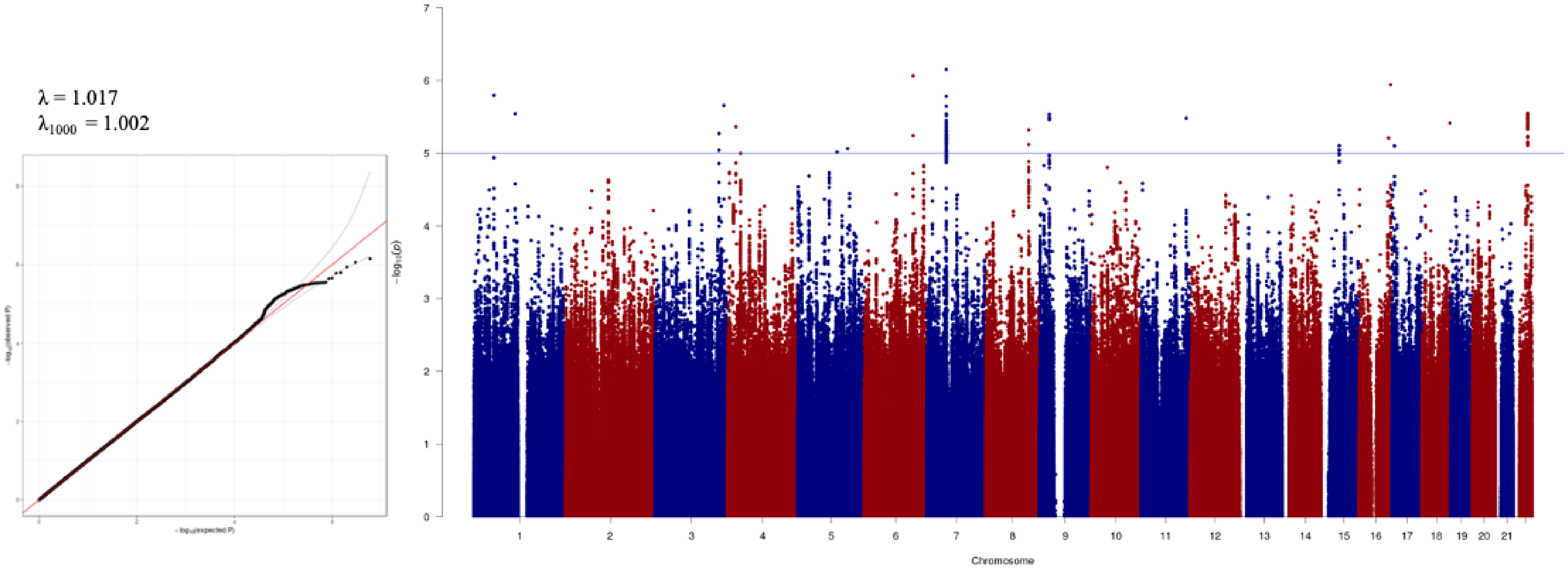


### Supplementary Figure 4. Manhattan plot and QQ plot with depression in individuals of South Asian ancestry.

The y-axes show the −log10*P* values of the association between each single-nucleotide variant and the outcome. The x-axes show the chromosomal position (GRCh37). The red line represents the genome-wide significance threshold of 5 x $10^{-8}$ and the blue line, $10^{-5}$.

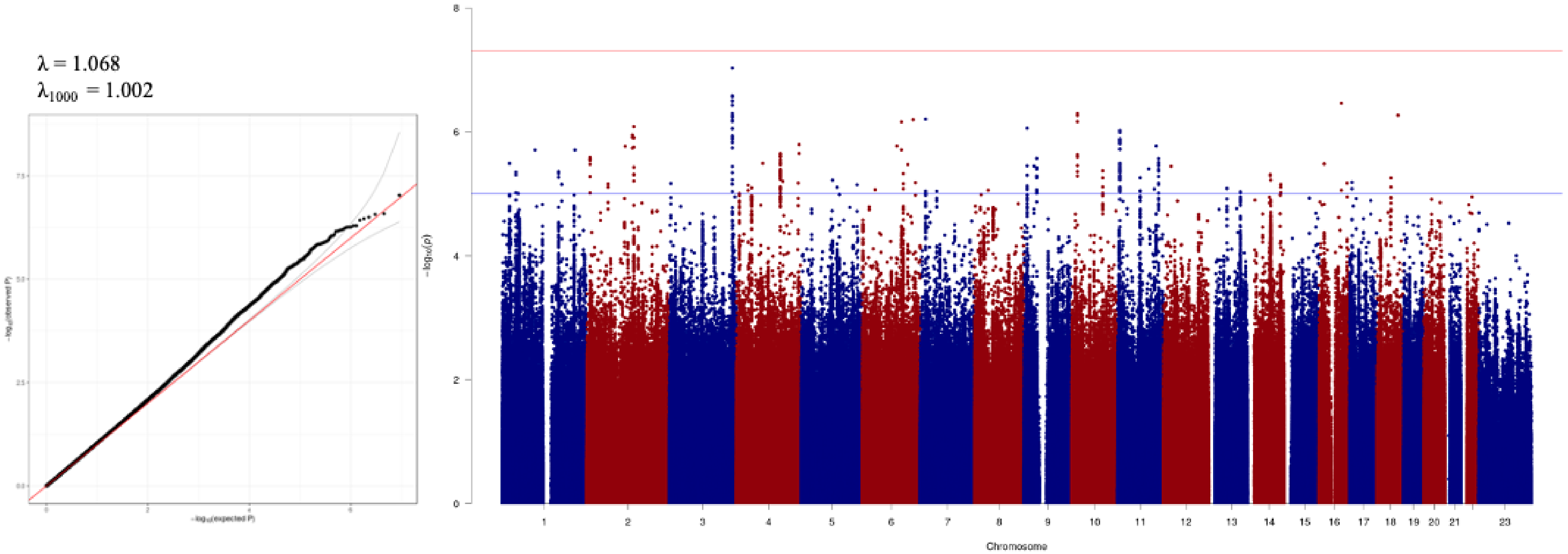


### Supplementary Figure 5. Manhattan plot and QQ plot with depression in individuals of East Asian ancestry.

The y-axes show the −log10*P* values of the association between each single-nucleotide variant and the outcome. The x-axes show the chromosomal position (GRCh37). The red line represents the genome-wide significance threshold of $5 \times 10^{-8}$ and the blue line, $10^{-5}$.

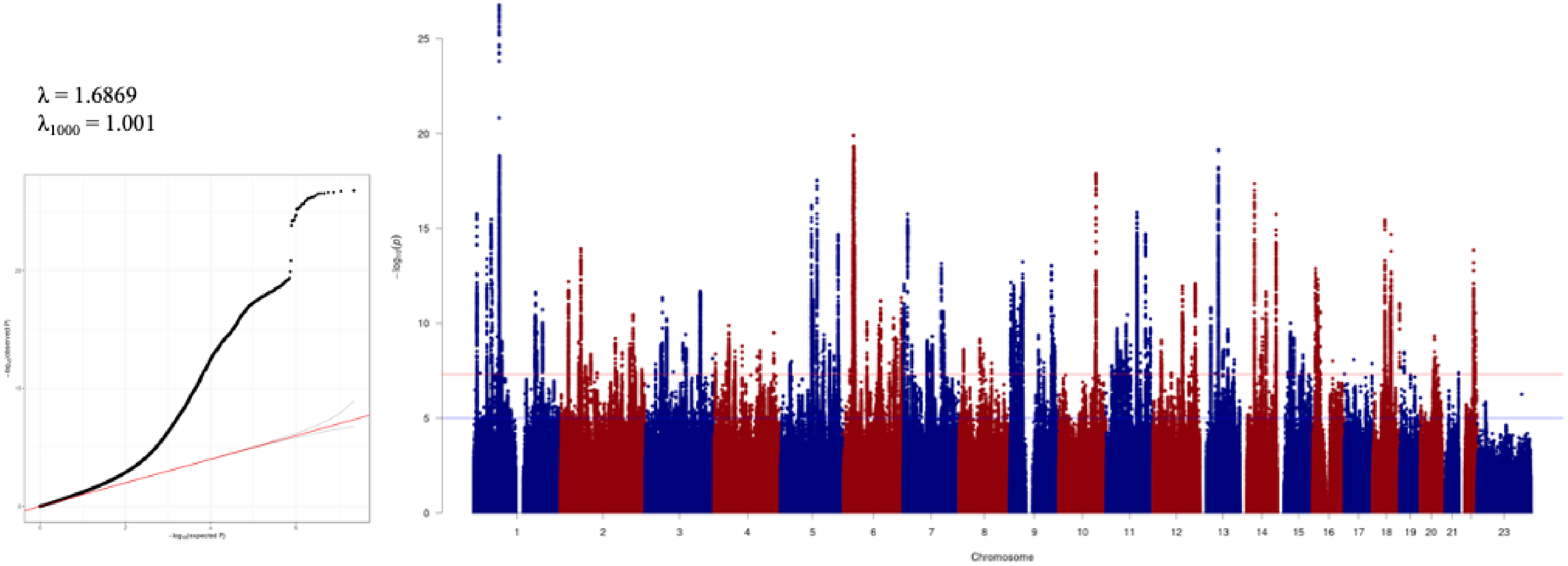


Supplementary Figure 6. Manhattan plot and QQ plot with depression in multi ancestry meta-analysis.

The y-axes show the −log10*P* values of the association between each single-nucleotide variant and the outcome. The x-axes show the chromosomal position (GRCh37). The red line represents the genome-wide significance threshold of $5 \times 10^{-8}$ and the blue line, $10^{-5}$. Association *P* values have been adjusted by the LDSC intercept of 1.0185.

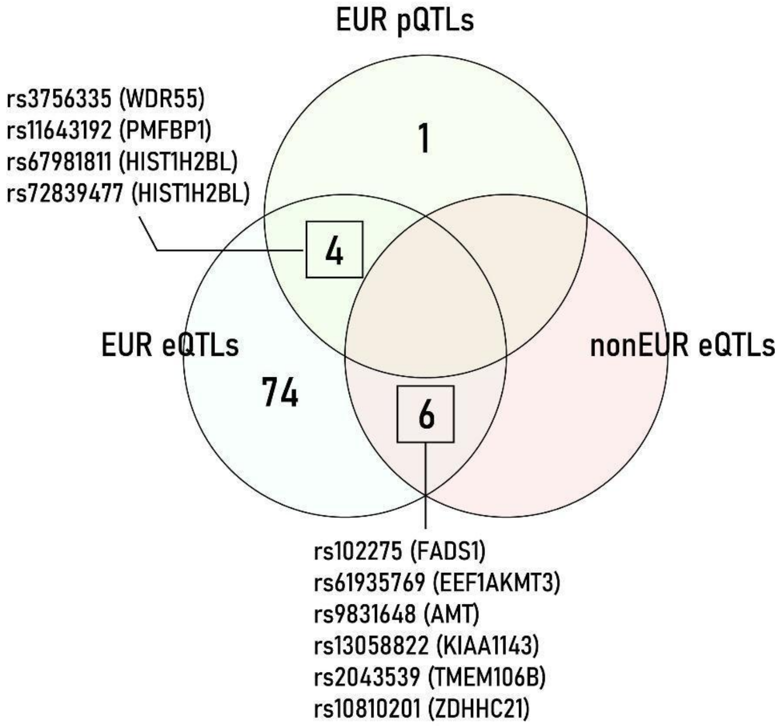


Supplementary Figure 7. Transferability SNPs mapped as QTLs from brain and blood tissue.

We show distribution of the 85 SNPs that are QTLs for brain and blood tissue for European and non-European studies.
