## Supplementary Material for "Multi-ancestry GWAS of major depression aids locus discovery, fine-mapping, gene prioritisation, and causal inference"

### Study Descriptions

A description of the following studies is available in our previous manuscript[1]: China Kadoorie Biobank (CKB), China, Oxford and Virginia Commonwealth University Experimental Research on Genetic Epidemiology cohort (CONVERGE), 23andMe cohort, Taiwan Major Depressive Disorder Study, Women's Health Initiative, Intern Health Study, UK Biobank, Army Study to Assess Risk and Resilience in Service members (Army STARRS), and BioMe.

Additional notes for the 23andMe cohort: Participants provided informed consent and participated in the research online, under a protocol approved by the external AAHRPP-accredited IRB, Ethical & Independent Review Services (E&I Review). Participants were included in the analysis on the basis of consent status as checked at the time data analyses were initiated.

The full GWAS summary statistics for the 23andMe discovery data set will be made available through 23andMe to qualified researchers under an agreement with 23andMe that protects the privacy of the 23andMe participants. Datasets will be made available at no cost for academic use. Please visit https://research.23andme.com/collaborate/#dataset-access/ for more information and to apply to access the data.

#### Genes & Health

Genes & Health (GH) is a community based, long-term study of health and disease in British Bangladeshi and British Pakistani people in east London[2]. GH uses a population-based study design, as well as incorporated genomics analysis, electronic health record (EHR) data, and targeted recall-by-genotype (RbG) studies[2]. As of February 2020 GH had 38 899 volunteers, and by 2023 it aims to reach 100,000 volunteers participating in the study.

The GH study design is divided into four stages:

**Stage 1: population-wide recruitment** - including volunteer questionnaire, electronic health record linkage, and a DNA for genotyping and exome sequencing

Bangladeshi and British Pakistani individuals (16 and over) living in or working in east London are invited to take part. Participants are recruited at mosques, libraries, GP surgeries and outpatient clinics by bilingual health researchers. Stage 1 participants all give consent to lifelong EHR linkage and donate saliva samples for genetic tests[2]. DNA was extracted from the Oragene (DNA Genotek) saliva system and stored from all Stage 1 volunteers. By late 2019, 50 000 samples from stage 1 volunteers were genotyped on the Illumina Infinium Global Screening Array v3.0 (with an additional 46 662 Multi-Disease variants)[2].

Data in volunteer questionnaires and EHR data was checked for data concordance and data with >99% concordance for gender and year of birth was retained. Data that could not be resolved with manual checking or data with clear data entry errors were removed[2].

**Stage 2: targeted recruitment** - recall-by-genotype studies
GH can invite participants four times per year for more detailed study visits, as part of Stage 2 study design, for clinical assessment and collection of biological samples. These visits and data collection are subject to further ethics approval, volunteer acceptability and community advisory group approval[2].

**Stage 3: Phenotype-focused studies** (from 2019 onwards)

**Stage 4: Therapeutic and preventative studies** (from 2019 onwards)

GH is aiming to genotype and high-depth exome sequence up to 100 000 volunteer samples by 2023[2].

### Biobank Japan

BioBank Japan (BBJ) is a prospective biobank that collected DNA and serum samples from 12 medical institutions in Japan between 2003 and 2008 and recruited approximately 200,000 participants[34].

Participants were recruited on the basis of having a disease diagnosis of at least 1 out of 47 target diseases[5]. All study participants provided written informed consent. The mean age of participants at recruitment was 63.0 years of age, and 46.3% of the participants were female. Participants were mainly of Japanese ancestry[4]. Data on medical history, drug prescription reports and biomarkers have also been collected [4].

**Initial Quality control of participants and genotypes in the initial BBJ**

Genotyping of participants was performed using Illumina HumanOmniExpressExome BeadChip or a combination of the Illumina HumanOmniExpress and HumanExome BeadChip[6]. Variants with the following criteria were excluded: call rate <99%, *P* value for Hardy–Weinberg equilibrium $<1.0 \times 10^{-6}$, and number of heterozygotes <5[6]. In the GWAS, Eagle[7] was used for haplotype phasing without an external reference panel and Minimac3[8] was used for imputation. Variants with an Rsq ≥0.3 were used in the association analysis[6].

**Further quality control in the 220 deep-phenotype GWAS data, used directly in the current study and conducted in the BBJ**

A total of 178,726 participants of East Asian ancestry were analysed (participants of the BBJ). The genotype data were further imputed with 1000 Genomes Project Phase 3 version 5 genotype data ($n = 2{,}504$) and Japanese whole-genome sequencing data ($n = 1{,}037$) using Minimac3 software[4]. Variants with an imputation quality of $Rsq < 0.7$ were excluded, resulting in 13,530,797 variants analysed in total [4].

GWASs for binary traits (disease endpoints and medication usage) were performed using a generalised linear mixed model implemented in SAIGE (v.0.37), adjusting for age, $age^2$, sex, age × sex, $age^2$ × sex and the top 20 PCs[4].

GWASs for quantitative traits (biomarkers) were performed using a linear mixed model implemented in BOLT-LMM (v.2.3.4), adjusting for the same covariates as in the binary traits above[4]. The inclusion of covariates for sex-specific diseases are described in detail elsewhere[4].

#### Million Veteran Program

The Million Veteran Program (MVP) is one of the world's largest programs on genetics and health, containing genetic data of over 870,000 veteran partners since launching the program in 2011. MVP is also one of the most diverse cohorts in the world, collecting data from participants across the United States of America[9]. The MVP is an observational cohort study combined with an electronic health record system, as well as a US Department of Veterans Affairs based mega-biobank.

<u>Data are being collected from participants in the format of:</u>
(1) questionnaires (a baseline survey on demographics, health status, medical history and a lifestyle survey on sleep, exercise, diet, and sense of wellbeing),
(2) through accessing electronic health records, and
(3) through a provided blood sample for genomic testing[10].

The study population is defined as active users of the Veteran Health Administration (VHA, large health care system) who are able to provide informed consent. 92.0% of the participants are male, between 50 and 69 years (55.0%). Self-reported race of 77.2% of the participants is white, and 13.5% is African-American[10]. Peripheral blood samples are collected by phlebotomists and processed plasma, buffy coat and DNA are stored in nitrogen freezers at The Department of Veterans Affairs Central Biorepository in Boston [10]. Genotyping (of approximately 459,777 samples) was performed using Affymetrix Axiom Biobank Array, with approximately 723K markers[11]. Sample quality control is described in detail in Hunter-Zinck et al.[11]

For the current study, we have used summary statistics from individuals of African and East Asian ancestry from MVP.

#### Australian Genetics of Depression Study (AGDS)

The Australian Genetics of Depression Study (AGDS) is one of the largest cohorts for studying genetic and psychosocial risk factors for depression[12]. For this study, more than 21,000 participants were recruited through two approaches: (1) through a nationwide recruitment based on pharmaceutical prescription history in the last 4.5 years and (2) through the Australian Department of Human Services and a media campaign (75% female, overall average age 43 years±15 years). Participants were asked to complete a self-reported online questionnaire assessing psychiatric history, clinical depression (using the Composite Interview Diagnostic Interview Short Form), responses to commonly prescribed antidepressants as well as voluntary questions assessing a range of traits relevant to psychopathology.

Participant enrolment was through the study website (https://www.geneticsofdepression.org.au/) hosted on a secure server (QIMR Berghofer). The website provided an information sheet about the study as well as a consent form for participants. Participants wishing to take part in the study were requested to sign the study informed consent and were asked to provide their name, age, and contact details. Participants' data were stored securely on the QIMR server[12]. To complete the questionnaire, participants were provided with a unique link to the questionnaires hosted on the Qualtrics website.

In addition to informed study consent, participants were asked to consent to provide access to their list of Medicare and Pharmaceutical Benefits Scheme (PBS) records for the previous 4.5 years. Consent was given by approximately 75% of the participants. Participants who did not consent to provide access to these records were still eligible to enrol to the study (around 25%).

Additionally, DNA samples were also collected from 75% of participants who agreed to provide their saliva samples via postal saliva kit. Saliva was self-collected by each consenting participant using an Isohelix GeneFix GFX-02 2 mL saliva collector, and a sample tube as well as a signed consent form specific to the treatment of genetic information were returned by post. DNA was extracted from the saliva sample and stored in freezers[12].

An additional clinical trial cohort of 214 individuals (66% female; mean age = 51.3, SD = 12.5, range = 21–79; mean BMI = 27.7, SD = 5.6) from a mental health cohort from Deakin University's IMPACT Institute was also included in the study, in order to increase the sample size. All blood samples in this cohort were from individuals who provided signed informed consent for further unspecified use of their samples[12].

Genotyping of participants from the study cohorts was done using the Illumina Global Screening Array V.2.0. for 15,792 participants as of November 2021[12]. The AGDS GWAS only consisted of participants who met the DSM-5 criteria for MDD and had not participated in previous studies contributing to Psychiatric Genomics Consortium (PGC) https://pubmed.ncbi.nlm.nih.gov/34924174/ The final sample size for the GWAS was 13,104 cases from AGDS and 214 cases from the Deakin University samples. The GWAS was conducted using the SAIGE v. 0.39 [13]. Results were filtered on a minor allele frequency > 0.01 and $R^2$ INFO score > 0.6 [12].

#### Genetic Epidemiology Research on Adult Health and Aging (GERA)

Individual data of the GERA study was accessed via dbGaP under project ID of 18933. The GERA study is a sub-study of the Kaiser Permanente (KP) Research Program on Genes, Environment, and Health (RPGEH), which consisted of 103,006 adult members of Kaiser Permanente North California (KPNC), ranging from 18 to 100 years at enrollment. The study design for RPGEH and GERA has been described in detail elsewhere[14]. Briefly, adult members of KPNC were asked to complete a mailed survey, the respondents of which completed a broad written consent and provided a saliva sample for DNA extraction. GERA participants were also asked to self-report their race, ethnicity, nationality and religion in order to maximize the diversity of the cohort. As a result, the final cohort was formed by 19% non-European individuals and 81% randomly drawn non-Hispanic white individuals. Phenotypes on demographic and behavioural factors (e.g., gender, marital status, education, smoking, alcohol consumption) were derived from the RPGEH surveys. Data on the occurrence of health conditions in participants in the GERA Cohort had been derived from summarising ICD-9 coded diagnoses in Kaiser Permanente's electronic medical records. An algorithm that aggregates specific ICD-9 codes into appropriate diagnostic groups for selected conditions was applied to outpatient and inpatient databases[15]. The criterion for counting a condition as "present" for a participant is the occurrence of two or more diagnoses within a diagnostic category occurring on separate days.

DNA was extracted from participants' saliva samples. The DNA extraction, genotyping, imputation, and relevant quality control procedures for the GERA cohort had been described elsewhere[16,17]. In short, GERA individuals were genotyped separately on four ethnic-specific arrays (Affymetrix Axiom) according to their self-reported ancestry. There were 84,430 participants assayed on the non-Hispanic European array, 8,043 on the East Asian array, 5,779 on the African array and 11,585 on the Hispanic/Latino array, respectively. The quality control for samples included DishQC score (DQC) of no less than 0.82 and sample call rate of no less than 97%. DishQC is a measure of the contrast between the AT and GC signals assayed in non polymorphic test sequences (Affymetrix White Paper). It provides a type of signal-to-noise figure of merit that is well correlated with sample call rate, allowing for prediction of successful samples. Afterwards, genotypes were imputed with the 1000 Genomes (October 2014 release) as a reference panel, implemented per array by IMPUTE2 v2.3.1[18,19].

The genetic ancestry based on the genotype imputation data was calculated by Graf-pop [20]. In total, we have got 226 MD cases and 3722 controls for East Asian ancestry, 206 cases and 1598 controls for African ancestry, 415 cases and 2808 controls of Hispanic/Latinx ancestry. GWAS were implemented for these three ancestry groups separately by PLINK2 on the imputed doage data, adjusting for sex, age and the first 10 PCs.

#### Jackson Heart Study (JHS)

Data of the Jackson Heart Study (JHS) was accessed via dbGaP under project ID of 18933. The participants with available depression phenotypes from a sub-study of JHS with genotyped dataset were analysed for the current study (dbGaP study accession: phs001356.v1.p2). Study designs for the JHS study have been described elsewhere[21]. In short, the JHS is a large, community-based, observational study, which recruited 5,301 participants from among the non-institutionalized African-American adults from urban and rural areas of the three counties (Hinds, Madison, and Rankin) that make up the Jackson, MS, metropolitan statistical area (MSA). Participants provided extensive medical and social history and had an array of physical and biochemical measurements and diagnostic procedures during a baseline examination (2000-2004) and two follow-up examinations (2005-2008 and 2009-2012). For current depression, cases were identified by a 20-item CES-D score of 16 or greater.

Study participants were genotyped by the Illumina Human Exome BeadChip v1.1 array and then imputed to the 1000 Genomes Phase 3 African reference panel. Samples with genotyping call rate of less than 95% and variants which were successfully genotyped in less than 95% of samples were excluded. Relatedness coefficients were calculated by KING[22]. Related individuals up to 2nd degree relatedness were randomly excluded (kinship > 0.0884).

A total of 299 depression cases and 990 controls were included in our GWA for the JHS study. Logistic regressions adjusting for age, gender, recruitment type and first 20 PCs were implemented in PLINK2. Following the analyses, variants with imputation R squared of less than 0.7 or a minor allele count of less than 50 were excluded.

#### The Drakenstein Child Health Study (DCHS)

The DCHS is a population-based birth cohort study in the Drakenstein area in Paarl, a peri-urban area 60 km outside Cape Town, South Africa. Data collection occurred at two clinics (maternal data) as well as at a central hospital (newborn outcomes) in the Drakenstein area between March 2012 to March 2015. Participants were enrolled in the DCHS at 20 to 28 weeks' gestation upon presenting for antenatal booking and followed longitudinally throughout pregnancy until at least five years postnatally. Maternal, paternal and child health are investigated through longitudinal measurements of risk factors in seven areas (environmental, infectious, nutritional, genetic, psychosocial, maternal and immunological) that may impact on child health[23,24]. More detailed information on the DCHS can be found on the study website[25]. Exclusion criteria for the DCHS were minimal in order to maximise generalisability, and focused primarily on those individuals who did not live in the region (and thus could not be readily followed up) or those who were intending to move out of the district within the first year. Exclusion criteria for cases and controls consisted of women who had stillbirths, infant deaths, gave birth to twins/triplets, were diagnosed with lifetime bipolar disorder or psychosis[26].

DNA was extracted from whole blood using the QIAsymphony DSP DNA Midi kit and protocol (Qiagen, Hilden, Germany). Genome-wide SNP genotyping was conducted using either the

Infinium PsychArray or Global Screening Array-24 BeadChip (Illumina). Standard quality control of the genome-wide data was performed using PLINK removing individuals with >5% missing data and removing one in each pair of related individuals with an IBD proportion >0.12 (indicating cousins or a closer relation). The DCHS researchers removed SNPs with call rates <95%, MAF <0.05 and deviation from Hardy–Weinberg proportions ($P < 1 \times 10^{-6}$ in controls and $P < 1 \times 10^{-10}$ in cases). To evaluate population stratification, PC eigenvectors of the genetic relationship matrix were calculated by using about 50 000 independent SNPs. SNPs in LD (r2=0.075) were excluded to calculate PCs. Dimensional plots of the PCs were also used to remove outliers. The Faculty of Health Sciences human research ethics committee of the University of Cape Town (UCT) approved this study.

We acquired individual-level genotype imputed data from the DCHS researchers. Depression cases were defined by BDI-II score. Controls were defined with both BDI-II score and EPDS score. There were 139 cases and 346 controls in our analysis. Logistic regressions were run by PLINK2 adjusting for the first 20 PCs, recruitment site, age of enrolment for controls or age of assessment for cases.

#### Vanderbilt University Biobank

The Vanderbilt University Medical Center's (VUMC, Nashville, USA) Biobank (BioVU) includes more than 285,000 patients seen at the VUMC, whose DNA was extracted (from whole blood) and linked to de-identified electronic health records data spanning 1990–2017[27]. A detailed consent form, including information on policies, data sharing and privacy is provided to patients seen at the clinic at VUMC. Signed informed consent of the patient is required to deposit new blood samples left over from clinical care to the BioVU Biobank. BioVU is overseen and approved by the VUMC Institutional Review Board.

**Genotyping and Quality control**

Genotyping was performed on a subset of BioVU patients ($n = 24{,}262$) using the Illumina MEGA$^{EX}$ platform, which contains more than two million markers. Quality control was done as described in Ruderfer et al. 2020[28]. Briefly, samples with greater than 2% missingness or abnormal heterozygosity were removed. Variants with greater than 2% missingness or Hardy-Weinberg equilibrium $p$-value $< 5 \times 10^{-5}$ were excluded. SNPs with minor allele frequency less than 2% and SNPs not genotyped in HapMap2 were also excluded. Imputation was performed using the pre-phasing/imputation stepwise approach in IMPUTE4 / SHAPEIT, using 1000 genomes phase I reference panel. Variants with INFO <0.3 were excluded [28].

A subset of SNPs in linkage disequilibrium was used to calculate relatedness and principal components of ancestry using multidimensional scaling in PLINK v1.9[29]. One individual from pairs of highly related individuals (pihat > 0.1) were randomly excluded. Samples were genotyped in five batches, and variants were removed if allele frequencies differed significantly ($P < 5 \times 10^{-5}$) between any batch and the rest of the sample[29]. Finally, multiallelic and structural variants were filtered, dosage data was converted to hard genotype calls, and variants with certainty <0.9 or INFO < 0.95 were excluded, resulting in 5,218,407 high quality SNPs across the autosomes[28,29].

#### Hispanic Community Health Study/Study of Latinos (HCHS/SOL)

HCHS/SOL is a prospective, multicenter, population-based cohort study of Hispanic/Latinx adults in the United States. It recruited 16,000 Hispanic/Latinox participants from Bronx, Chicago, Miami and San Diego under a two-stage area probability sampling[30]. All participants went through thorough baseline examination, which lasts for 7 hours on average. Signed informed consent was obtained when participants arrived at assessment centres, followed by fasting state measurements (e.g. anthropometry, phlebotomy, 2-hour glucose load) and several other measurements (e.g., EGC, seated blood pressure). Participants were also administered with a questionnaire collecting their socio-demographic status, medical history, substance use, wellbeing, etc.[31]. The HCHS/SOL study was approved by institutional review boards at participating centres, and written informed consent was obtained from all participants.

DNA extracted from blood was genotyped on an Illumina custom array, SOL HCHS Custom 15041502 B3, consisting of the Illumina Omni 2.5M array (HumanOmni2.5-8v1-1) and ~150,000 custom SNPs selected to include ancestry-informative markers, variants characteristic of Amerindian populations, previously identified GWAS hits, and other candidate-gene polymorphisms. Genotype imputation was performed with the 1000 Genomes Project phase 1 reference panel implemented by SNAPEIT2 and IMPUTE2[32].

Depressive symptoms were assessed during the baseline assessment with the Andresen version of the 10-item Center for Epidemiology Studies of Depression Scale (CES-D-10), which reflected core symptoms of depression in the past week[33]. The CES-D-10 scores were curated into a binary phenotype, where participants with scores of no less than 10 were defined as cases (N = 3,979) and those with scores of 6 or below were defined as controls (N = 6,499). Mixed-effect model logistic regressions adjusting for log of sampling weight, recruiting centre age, sex, highest education attained, genetic subgroup and the first 5 PCs were conducted by GENESIS[33,34].

#### Detroit Neighborhood Health Study (DNHS)

The DNHS is an ongoing, longitudinal epidemiologic study investigating correlates of PTSD and other mental disorders in the city of Detroit. It recruited adults (18 years or above) from the Detroit population. Initially, 1,547 households were randomly drawn from the city of Detroit under a probability sample. Then an interview was conducted with one random individual from each household. Participants underwent a 40-minute assessment consisting of questions on socio-demographic characteristics, major depressive disorder (MDD), and generalized anxiety disorder (GAD)[35]. MD was scored using the Patient Health Questionnaire

(PHQ-9)[36] and DSM-IV criteria[37]. The Institutional Review Board at the University of Michigan and the University of North Carolina Chapel Hill approved this study.

DNA for GWAS analysis was isolated from peripheral blood or saliva. Study participants were genotyped with Illumina HumanOmniExpress array and imputation was conducted based on the 1000 Genomes phase 3 data. Relatedness was estimated using the IBS function in PLINK 1.9. From each pair with relatedness $\pi\^{} > 0.2$, one individual was removed from further analysis, retaining cases where possible. Principal components were calculated based on the smartPCA algorithm in EIGENSTRAT[38].

A total of 58 cases, which were defined at the baseline visit, and 436 controls of African ancestry were included in our MD GWAS for the DNHS cohort. Individual level genotype imputed data and phenotype data were shared with us. Logistic regressions were implemented by PLINK2 with imputed dosage data, adjusting for sex, age at baseline assessment, and the first 20 PCs.

#### Prevention Intervention Research Center (PIRC) 1st Generation Trial

This study was a trial designed and conducted by the Prevention Intervention Research Center at Johns Hopkins University. A total of 2,311 youth entering first grade in 1985 or 1986 in 19 primary public schools, which were selected from five areas to represent the socio-demographic diversity in the northeastern quadrant of Baltimore in 1985, were recruited[39]. The trial was initially aimed at assessing the immediate effects of two universal, first-grade preventive interventions (i.e., classroom-centered intervention vs family-school partnership intervention) on the proximal targets of poor achievements, concentration problems, aggression and shy behaviours, which were known as early risk behaviours for later substance use/abuse, affective disorder and conduct disorder[40]. Twenty five years later, 65% of the surviving cohort (n=1,434) participated in a follow-up interview that inquired about their general and mental health, including alcohol, tobacco, and other drug involvement. Lifetime history of major depressive episode was measured using the

Diagnostic Interview Schedule-III-R (DIS-III-R). The study protocol was approved by the institutional review board for protection of human subjects at Johns Hopkins University [39].

Participants were asked about their willingness to donate a blood sample. If unwilling and/or unable to donate blood, they were then asked if they would donate a saliva sample instead. DNA was extracted from blood or saliva samples, then quantitated and genotyped using Affymetrix 6.0 microarray (Santa Clara, CA, USA)[41]. Genotypes were imputed to the TopMed using the Michigan Imputation Server[42]. The resulting variants imputed with an INFO score of less than 0.8 were removed. Genome-wide logistic regressions were conducted by R (version 3.6.1) for 52 cases and 547 controls of African ancestry, adjusting for participant age, sex, intervention status (control vs exposure to intervention) and the first 20 PCs.

#### Mexican Adolescent Mental Health Survey (MAMHS)

MAMHS is a multistage probability survey of 3,005 adolescents aged 12-17 years residing in Mexico City. Interviews were conducted in the homes of the invited participants in 2005. Signed informed consents were obtained from parents or legal guardians, and assent of the participating adolescents were obtained as well[43]. Lifetime depression was defined by the MDE DSM IV criteria, as with the computer-assisted version of adolescent CIDI.

DNA was extracted from exfoliated oral cavity cells and afterwards genotyped with the Illumina Global Screening Array (GSA). Imputation was carried out using the Michigan Imputation server using Minimac4 and the 1000 g-phase-3-v5 (hg19) full reference panels [44]. Variants with imputation INFO of less than 0.7, minor allele frequency of no larger than 0.01, missingness no less than 0.05 or with Hardy-Weinburgh *P* value of no larger than $1\times10^{-6}$ were excluded. Participants with genotype missing rate of no less than 0.05 or heterozygosity outliers (> 3 standard deviation away from the mean) were excluded, leaving 105 cases and 996 controls of admixed Hispanic/Latinx ancestry for GWA analysis. Mixed-effect model logsitic regressions were implemented by SAIGE [13] adjusting for age, sex and the first 20 PCs.

#### Pregnancy Outcomes, Maternal and Infant Study (PrOMIS)

The PrOMIS cohort is a prospective cohort aimed at understanding the life course and intergenerational effects of interpersonal violence and other forms of trauma among Peruvian women. Between 2012 and 2015, participants were recruited from prenatal care clinics at the Instituto Nacional Materno Perinatal (INMP) in Lima, Peru. A structured questionnaire including maternal socio-demographic, lifestyle characteristics, medical and reproductive histories, and mental health symptoms was completed by an interview with trained research staff. Depression was assessed with the Patient Health Questionnaire (PHQ-9), which enquired about depressive symptoms for the 2-week period prior to the interview. The institutional review boards of the INMP and the Office of the Human Research Administration, Havard T.H. Chan School of Public Health approved all procedures used in the study[45,46].

Genotyping was conducted on the Illumina Multi-Ethnic Global Chip. Imputation was conducted with the 1000 Genomes phase 3 data[47]. GWAS quality control and imputation was performed to the published PGC procedures[38]. PC-related and PC-Air was employed in identifying and excluding related individuals, and calculating principal components[48]. There were 1,076 MD cases and 2,328 controls for the MD GWAS. GWA logistic regressions were conducted by PLINK, adjusting for the first 10 PCs.
